# Cluster Assembly Dynamics Drive Fidelity of Planar Cell Polarity Polarization

**DOI:** 10.1101/2024.10.21.619498

**Authors:** Silas Boye Nissen, Alexis T. Weiner, Kaye Suyama, Pablo Sanchez Bosch, Maiya Yu, Song Song, Yuan Gu, Alexander R. Dunn, Jeffrey D. Axelrod

## Abstract

In planar cell polarity (PCP) signaling, distinct molecular subcomplexes segregate to opposite sides of each cell, where they interact across intercellular junctions to form asymmetric clusters. Although proximal-distal asymmetry within PCP clusters is the defining feature of PCP signaling, the mechanism by which this asymmetry develops remains unclear. Here, we developed a method to count the number of monomers of core PCP proteins within individual clusters in live animals and used it to infer the underlying molecular dynamics of cluster assembly and polarization. Measurements over time and space in wild type and in strategically chosen mutants demonstrate that cluster assembly is required for polarization, and together with mathematical modeling provide evidence that clusters become increasingly asymmetric and correctly oriented as they increase in size. We propose that cluster assembly dynamics amplify weak and noisy inputs into a robust cellular output, in this case cell and tissue-level polarization.

## INTRODUCTION

The assembly of proteins into large, complex assemblies is a ubiquitous aspect of both signal transduction cascades and the protein complexes that mediate adhesion between neighboring cells^1^. Both classes of molecular assemblies, for example cadherin-mediated adherens junctions and the Wnt signalosome, have features in common with biological condensates such as large and variable sizes and complex stoichiometries^2^. In general, the functions of cluster formation in these and other systems are still incompletely understood. This knowledge gap reflects, in part, the experimental difficulties associated with probing how large protein assemblies form, and how their formation and dynamics underlie biological function.

Molecular counting provides one means of gaining insight into how large protein assemblies form and function. Inspired in part by existing theoretical frameworks^3^, previous researchers have used protein cluster size distributions to gain insight into cluster assembly dynamics and the resulting consequences for biological function. The E-cadherin cluster size distribution in the *Drosophila* embryonic epithelium was found to obey a power law distribution with a superimposed exponential decay that limits the largest sizes^4^. From this, it was inferred that cluster assembly is regulated by a combination of fusion and fission, along with the selective removal of large clusters. In another example, an exponential size distribution for clusters of the scaffolding protein PAR-3 was found in *C. elegans*^5^. Together with additional experimental evidence, this led to the proposal that two positive feedback mechanisms stabilize the asymmetric distribution of PAR-3^6,7^.

In planar cell polarity (PCP) signaling, six core components, Flamingo (Fmi)^8,9^, Frizzled (Fz)^10,11^, Van Gogh (Vang)^12,13^, Dishevelled (Dsh)^14,15^, Diego (Dgo)^16^, and Prickle (Pk)^17^ break cellular symmetry by segregating distinct subcomplexes to opposite sides of the cell^18^. The two subcomplexes communicate intercellularly, recruiting one subcomplex (Fmi-Fz-Dsh-Dgo) to one side of the cell junction, and the other subcomplex (Fmi-Vang-Pk) to the opposite side (Fig. 1A). In *Drosophila* wing cells, the two oppositely localized subcomplexes direct cytoskeletal regulators to orient the growth of pre-hairs^19,20,21^. In vertebrates, including humans, PCP is essential for a variety of developmental processes, for example neural tube closure and heart formation, and for several physiological and pathological processes such as wound healing and cancer invasion^18^.

**Figure 1:**
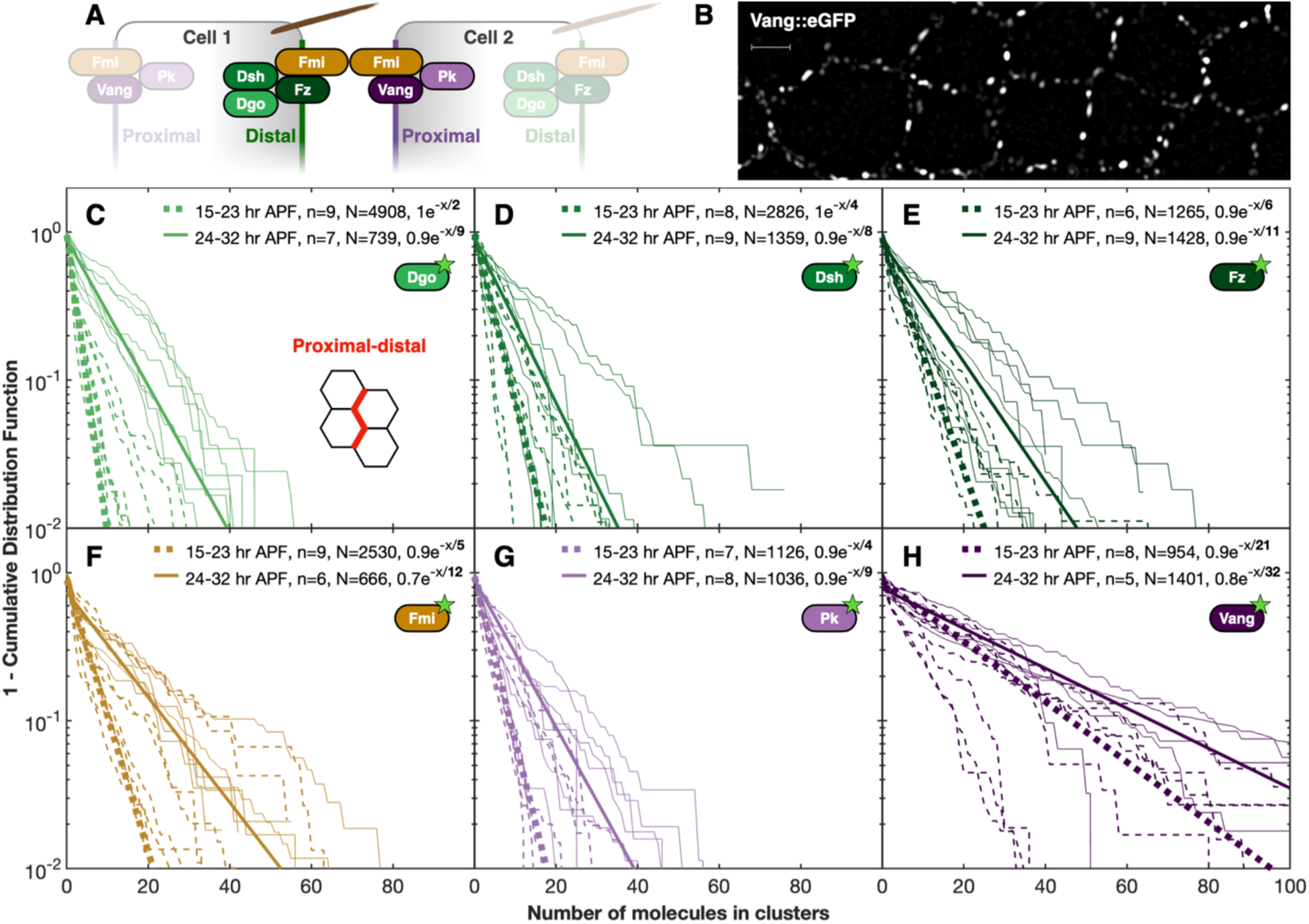
Numbers of core component molecules in PCP clusters follow single exponential distributions. **(A)** Cartoon of a PCP signaling cluster. The distal subcomplex consists of Diego (Dgo), Dishevelled (Dsh), and Frizzled (Fz). The proximal subcomplex consists of Prickle (Pk) and Van Gogh (Vang). Flamingo (Fmi) is in both subcomplexes and connects them across intercellular junctions. **(B)** High-resolution image of Vang::eGFP acquired with Total Internal Reflection Fluorescence (TIRF) Microscopy 29 hr after puparium formation (APF, see *Methods* for details). Vang assembles into clusters of varying sizes along cell-cell junctions. Scale = 2 μm. **(C-H)** Cluster size distributions of **(C)** Dgo, **(D)** Dsh, **(E)** Fz, **(F)** Fmi, **(G)** Pk, and **(H)** Vang (each tagged with GFP), grouped into an early age group (15-23 hr APF, dashed lines) and a late age group (24-32 hr APF, solid lines) gated on proximal-distal boundaries. Thin lines indicate individual samples (each from independent wings). Solid lines illustrate a single exponential fit through a median wing. *n* indicates the number of wings imaged, and *N* indicates the total number of clusters measured. Note the logarithmic *y*-axis. The average number of each component in clusters corresponds to the denominator of the exponent. See Figure S1 for anterior-posterior boundaries.

Construction of the core PCP complex Dgo-Dsh-Fz-Fmi=Fmi-Vang-Pk (Fig. 1A) that spans intercellular junctions (indicated by =) was proposed to involve the initial assembly of the transmembrane components into an asymmetric bridge comprising Fz-Fmi=Fmi-Vang^22^. Addition of the cytoplasmic components (Dgo, Dsh, and Pk) forms a complete, asymmetric complex (with asymmetry here defined as unequal amounts of Fz, Dgo, Dsh, Vang and/or Pk on the two sides of the complex). A critical element of this model is a preferential asymmetry, such that Fz-Fmi=Fmi-Vang is preferred over symmetric Fz-Fmi=Fmi-Fz or Vang-Fmi=Fmi-Vang^23^. While it is well established that asymmetric signaling is transmitted between cells by Fmi=Fmi bridges^22–26^, the basis for how this initial asymmetry is amplified and correctly oriented relative to the overall proximal-distal axis is not certain.

In confocal images, asymmetry appears to be greatest in junctional regions containing clusters of PCP complexes, which appear as bright puncta^27,28^. This observation suggests that ‘mutual exclusion’, wherein proximal and distal subcomplexes exclude each other from their respective domains^18^, is somehow associated with cluster formation. PCP clusters are inferred to contain all six core components, and cluster formation is likely mediated by several previously observed phenomena. Overexpression of Dsh or Pk results in the formation of huge puncta^16,29,30^, and in the absence of either protein, the size and stability of puncta are diminished^27,28^. Oligomerization may contribute to clustering, as the DIX domain in Dsh and a poorly characterized domain in Pk mediate oligomerization^31,32^. N-terminal Vang phosphorylation sites appear to modulate clustering^31,33,34^ and Fmi *cis*-interactions may promote clustering^35^. Clustering may be associated with stability: the localization of Fmi and Fz within large puncta is more stable than elsewhere, as judged by fluorescence recovery after photobleaching (FRAP)^28^. Surprisingly, the stoichiometry of complexes appears not to be fixed: semiquantitative imaging revealed that modest changes in expression levels of any cytoplasmic components altered their incorporation into complexes without changing the relative abundance of other components^36^.

Whether cluster formation in PCP signaling is functionally significant, and if so, in what way, is unknown. For example, cluster size might be related to asymmetry (defined as above), proper orientation (in the wing, defined as Fz, Dgo and Dsh accumulating on the distal side and Vang and Pk accumulating on the proximal side of cells), and/or to the ability to transduce a downstream response. We hypothesized that elucidating the PCP cluster size distribution would yield insight into the underlying functional importance of clusters in PCP signaling. To explore this hypothesis, we developed a method to quantify the number of monomers of the core PCP proteins within individual complexes, thereby defining distributions and relationships in various wild-type and mutant conditions, including in mutants that specifically impair clustering. We then captured these measurements, prior knowledge, and data from a companion manuscript (Weiner et al.^37^) to suggest a mathematical model of cluster formation and a mechanism for symmetry breaking. From these studies, we draw an overarching conclusion that growth of sufficiently large clusters is necessary for cell polarization; we suggest that the asymmetry of individual clusters and the probability of correct cluster orientation increase with size, and we propose that this is achieved by a cluster growth mechanism that constitutes a proofreading function that strengthens as a function of cluster size.

## RESULTS

### The core PCP signaling proteins assemble into clusters that follow a size-independent growth mechanism

We hypothesized that quantifying the distribution of PCP cluster sizes would yield insight into how clusters assemble, and that carefully defining cluster composition could provide insight into how symmetry breaking occurs. Among many possibilities for growth, clusters may grow to a fixed, uniform, cluster size distribution where the on- and off-rates equilibrate. Alternatively, clusters might undergo coarsening, continually fusing to result in bimodal cluster size distribution, or a combination of fusion and fission might yield a power law distribution^38^. Bistability, proposed as a central feature of PCP signaling^39^, might be encoded in a cooperative mechanism resulting in nonlinear cluster growth, *i.e.* positive feedback in the recruitment of individual components into clusters. None of these possibilities have been examined to our knowledge.

For quantification, we employed eGFP-tagged core PCP proteins previously validated to express and incorporate into clusters at levels similar to the endogenous proteins (except for the Fz::eGFP probe that is thought to incorporate roughly twice as efficiently as endogenous Fz; no correction for this effect is made in the present analyses)^36^. Imaging at high spatial resolution in live tissue by TIRF microscopy, we observe many more clusters of varying sizes than previously appreciated (Fig. 1B). To obtain cluster size distributions, we count the number of fluorophores in each cluster by identifying fluorophore steps in the bleaching trace (see Methods).

For each of the six core proteins, at proximal-distal (P-D) boundaries, the molecular size distributions we observed are well fit by single exponential functions (Fig. 1C-H; see Methods). These data are inconsistent with models that predict uniform cluster sizes, coarsening (fusion), or a combination of fission and fusion. Instead, exponential cluster size distributions are parsimoniously explained by cluster growth and decay rates independent of the cluster size, with decay rates necessarily higher than the corresponding growth rates to prevent runaway cluster growth^40^. Importantly, the exponential cluster size distributions indicate that if indeed bistability underlies PCP asymmetry, we see no strong evidence that it is a consequence of cooperative growth of cluster size.

### Clusters grow and cluster stoichiometry is maintained

To determine whether the cluster size distribution evolves as cellular asymmetry (the segregation of Fz, Dgo, Dsh, from Vang, Pk within a cell) increases, we measured the evolution of cluster stoichiometry and size over time. A wave of cell divisions and a dramatic reorganization of microtubules between 14 and 16 hr after puparium formation (APF)^41^ disturb PCP, allowing us to assay cluster composition as cellular polarity is consolidating following this disturbance. At P-D boundaries, the average number of each molecule within clusters grows between 15 and 32 hr APF (Fig. 2). In contrast, we see no or little growth of clusters at A-P boundaries that are orthogonal to the polarity axis, highlighting that cluster size growth is most pronounced along the axis of polarization (Fig. S1 and S2). Our results for P-D boundaries differ modestly from previous estimates that found no growth of any component except for a small growth of Vang between 20 and 28 hr APF^36^.

**Figure 2:**
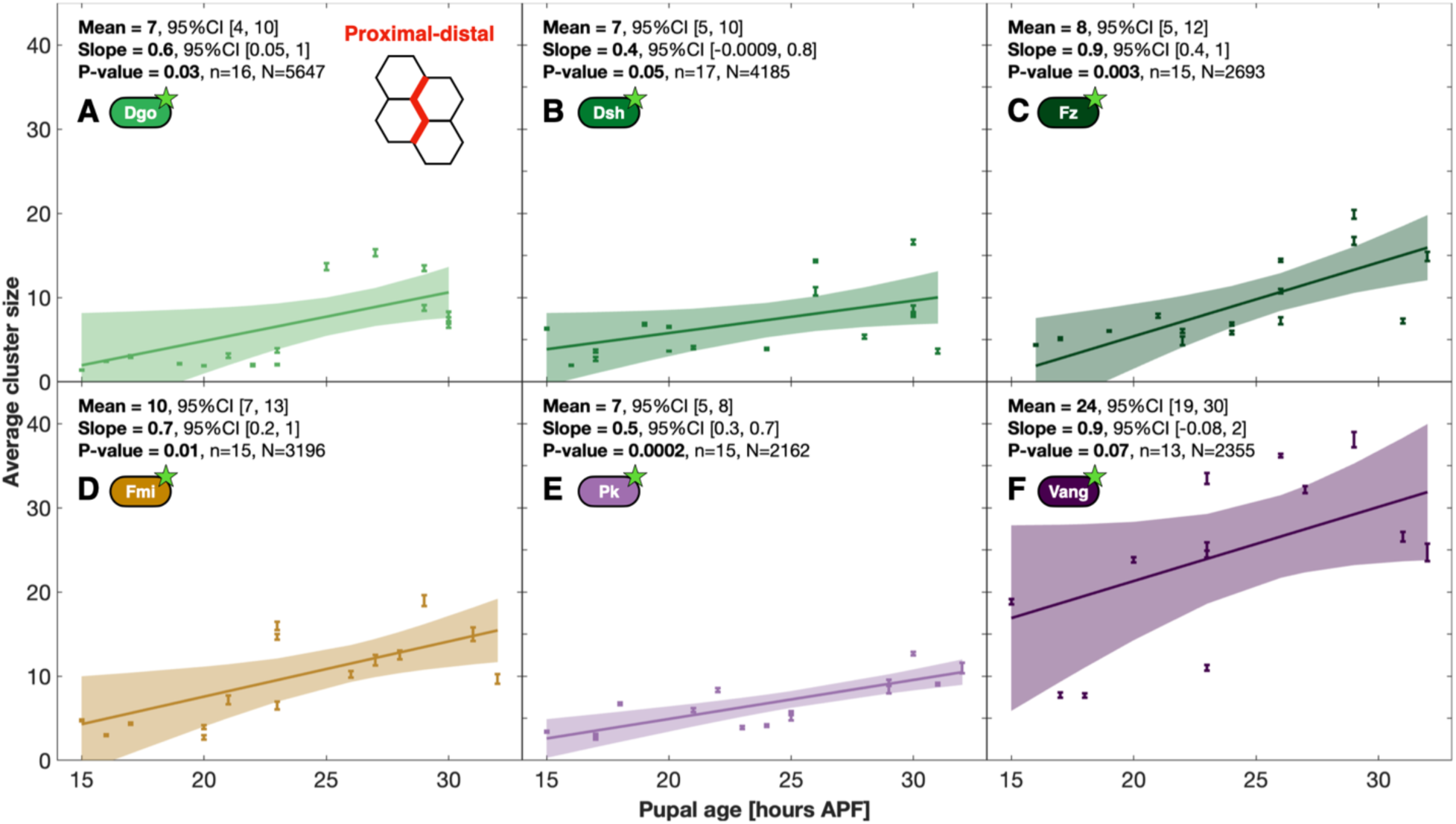
Stoichiometry is roughly maintained as PCP clusters grow. (A-F) The average number of monomers in clusters along proximal-distal boundaries as a function of the number of hr APF for **(A)** Dgo, **(B)** Dsh, **(C)** Fz, **(D)** Fmi, **(E)** Pk, and **(F)** Vang. Each dot represents a single-exponential fit through the cluster size distribution found in one sample and the error bar represents the standard error of the estimate (see Figure 1). For each core protein, a weighted linear regression model is shown through all samples (solid line) with 95% confidence interval (CI, shaded background). The mean indicates the average cluster size at 23.5 hr APF, and the slope represents the rate of increase in monomers/hour. The P-value tests the significance of the slope (see *Methods*). *n* is the number of biological samples, and *N* is the total number of clusters analyzed. For anterior-posterior boundaries, see Figure S2.

We estimated average numbers of components within individual clusters at 23.5 hr APF (Fig. 2). This result equates to an approximate stoichiometry of 3:1:1:1:1 for Vang, Fz, Dgo, Dsh, Pk respectively, with somewhat more than one Fmi relative to Fz, Dgo, Dsh and Pk. We note that this is an average, and it does not necessarily reflect the stoichiometry of individual clusters (see 2-color imaging below). The three-fold excess of Vang relative to Fz, Dgo, Dsh and Pk is consonant with the recently reported trimer structure of Vang^42,43^. We note, however, many clusters with one or two Vang monomers; the trimer structure of Vang may therefore not be obligate. The number of monomers of the six proteins in clusters grow at rates that maintain roughly the same stoichiometry during the 15 to 32 hr interval studied (Fig. 2).

The density of clusters remained roughly the same and independent of age (Fig. S3). We performed high resolution live imaging using TIRF and observe that while some larger clusters may persist for an hour or more, as previously observed^28^, many clusters appear to have lifetimes on the order of minutes; in addition, clusters show some mobility that decreases as their size increases (Movie S1 and Fig. S4). While we cannot rule out that some fission and fusion may occur, it appears that clusters undergo growth and decay for the most part independently of each other.

### Clonal analysis reveals that a population of distal Vang is present in PCP clusters

To determine how polarized each component is within individual clusters, we separately counted the number of molecules in proximal and distal subcomplexes using clonally expressed probes. All the resulting cluster size distributions again followed single-exponential distributions (Fig. S5). For Fmi, these data reveal equivalent size distributions (and numbers of clusters; Fig. S3) on the proximal and distal sides of junctions with an average of 5 Fmi monomers on each side at 23.5 hr APF (Fig. 3A,A’,A’’), consistent with a 1:1 *trans* homodimer configuration. The number of Fmi monomers on either side increases with an average net addition of one monomer every 2-3 hours.

**Figure 3:**
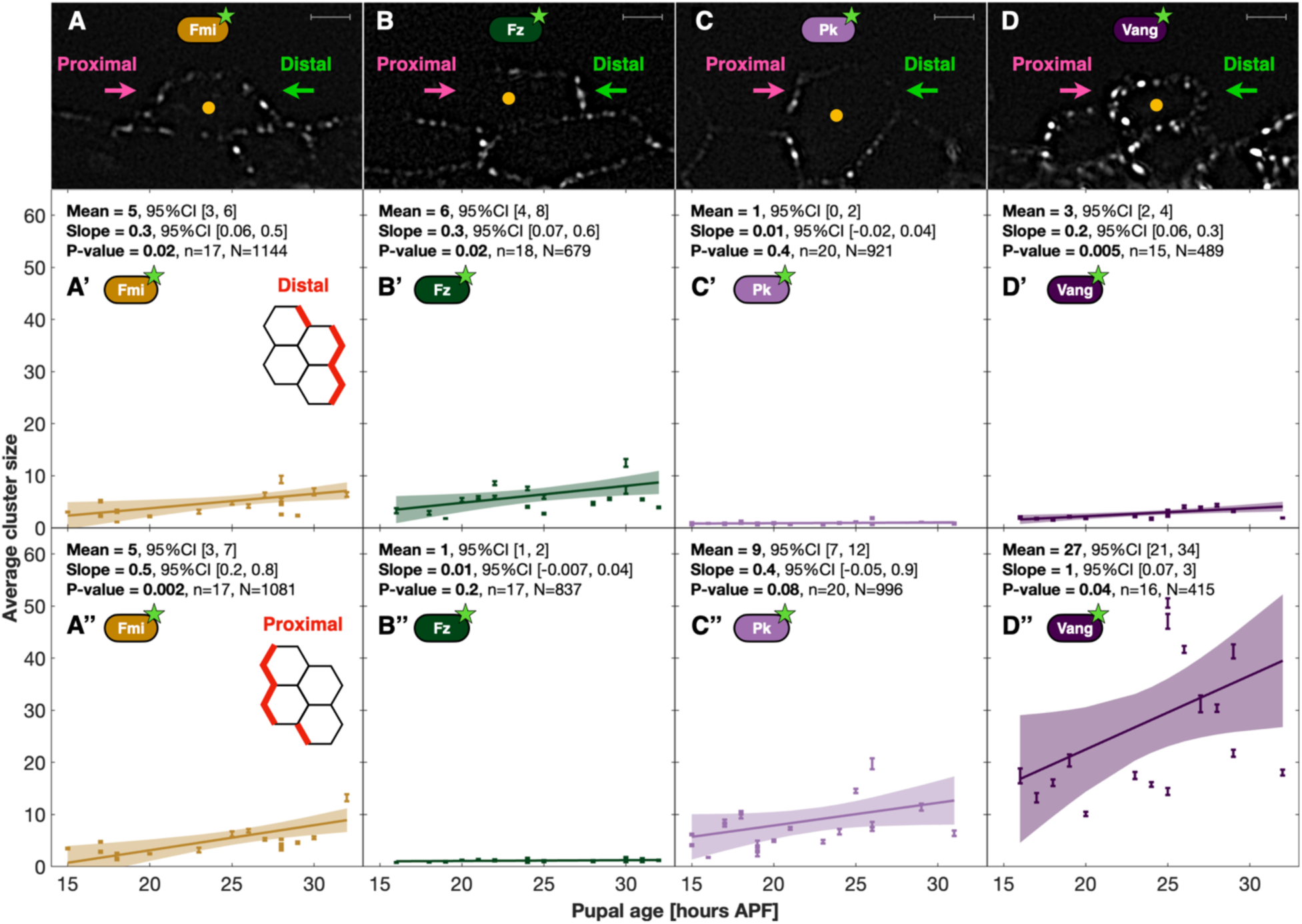
PCP protein polarization assayed by clonal analysis. (A-D) High-resolution sample images of **(A)** Fmi, 22 hr APF, **(B)** Fz, 22 hr APF, **(C)** Pk, 26 hr APF, (**D)** Vang, 26 hr APF, each expressed mosaically, all tagged with eGFP (see *Methods* for details). Orange dots indicate cell centers. All scale bars are 2 μm. The apparent paucity of Fz subcomplexes on the proximal side and Pk subcomplexes on the distal side is due to the choice of contrast in the images shown. In contrast, note the presence of visible distal Vang subcomplexes. **(A’-D’’)** The average number of monomers in subcomplexes based on single exponential fits through the cluster size distribution identified in individual samples (dots and error bars). **(A’-D’)** Subcomplexes along distal cell boundaries vs. along **(A’’-D’’)** proximal cell boundaries. (**A’, A’’)** Fmi. **(B’, B’’)** Fz. **(C’, C’’)** Pk. **(D’, D’’)** Vang. Note the presence of Fmi on both proximal and distal sides. Dots, error bars, Mean, Slope, P-value, *n* and *N* as in Figure 2. The underlying cluster size distributions are shown in Figure S5. For anterior-posterior boundaries, see Figure S2.

In contrast, Fz is highly polarized with most on the distal side and very little on the proximal side (Fig. 3B,B’,B’’). Fz was found in similar numbers on the distal side as Fmi, suggesting a roughly 1:1 stoichiometry between the two proteins on the distal side. The distal Fz population grows at the same speed as Fmi, with the net addition of one monomer every three hours, maintaining 1:1 stoichiometry (As noted above, the actual ratio in wild-type animals may be somewhat lower as the Fz probe incorporates more efficiently than endogenous Fz.) In contrast, we find a population of proximal subcomplexes that contain Fz at an average size of one monomer, that does not increase or decrease with age (Fig. S5B’).

Pk behaves essentially as a mirror image of Fz (Fig. 3C,C’,C’’). On the proximal side, clusters were of average size 9 at 23.5 hr APF and grow at a rate equivalent to that of distal Fz and Fmi. Distal Pk exists on average as single monomers and does not grow with age. We infer that many clusters have no Fz and Pk on the side where they are not dominant.

Proximal Vang exists in numbers larger than any other core protein (Fig. 3D,D’’): roughly three-fold higher than proximal Pk and five-fold higher than proximal Fmi. Proximal Vang grows at 3 monomers every 2-3 hours, maintaining the 1:3 Pk:Vang stoichiometry on the proximal side. Surprisingly, we find a population of Vang on the distal side with an average distal subcomplex containing 3 Vang monomers (Fig. 3D,3D’). The distal Vang population is smaller than both distal Fmi, and distal Fz, but 3-fold larger than distal Pk. This finding was not anticipated based on prevailing understanding, although it was evident but not commented upon in an earlier report^27^. We find a slow but significant growth in the distal Vang population with 1 monomer being added every 5 hours on average. Thus, growth in the number of Vang molecules occurs primarily on the proximal side, and at 30 hours, when the position of pre-hair growth is set, and cell polarization appears strongest, large clusters with large amounts of proximal Vang are present.

In sum, equal amounts of Fmi are present on the distal and proximal sides, whereas Fz, Pk, and Vang are roughly 90% polarized, with Vang present at an approximately three-fold excess over Pk.

### Clusters form in null mutants

Null mutants of each core protein produce substantial polarity defects and inhibit the asymmetric localization of the remaining core proteins, indicating that PCP is profoundly compromised at a functional level. Multiple interactions between core components have been reported^18^, and data suggest that several core proteins can oligomerize^31–35,44,45^, consistent with the possibility that clustering might not require the presence of the full complement of components. However, the extent to which clusters might form in the absence of any individual core PCP protein has not been quantitatively assessed.

We used molecular counting to determine how cluster assembly is altered in the absence of individual core proteins. For the loss-of-function conditions examined (*fz, vang, pk*), we observe clusters (Fig. S6). In *fz* mutants, Fmi forms clusters of similar size to wild-type at early time points, suggesting that Fmi incorporated into PCP clusters does not depend on Fz (Fig. 4A-A’’). Both Vang and Pk are recruited into clusters at levels comparable to wild-type at early time points (Fig. 4B-C’’), but the number of Vang monomers in clusters fails to grow with pupal age. In consequence, we find substantially reduced numbers of Vang monomers in *fz* mutant clusters at 32 hrs APF. This dependency suggests that a Fz-dependent signal from the distal side increases the recruitment rate of proximal Vang but is not required for its baseline recruitment (We show below that this signal depends not only on Fz but also on Dsh.) Quantifying *vang* or *pk* mutants likewise revealed that the number of Fz molecules within early clusters does not depend on the presence of Vang or Pk; however, growth of Fz levels depends on Vang, suggesting that a Vang-dependent signal from the proximal side likewise increases the recruitment rate of distal Fz (Fig. 4D-E’’; see also ref. ^46^).

**Figure 4:**
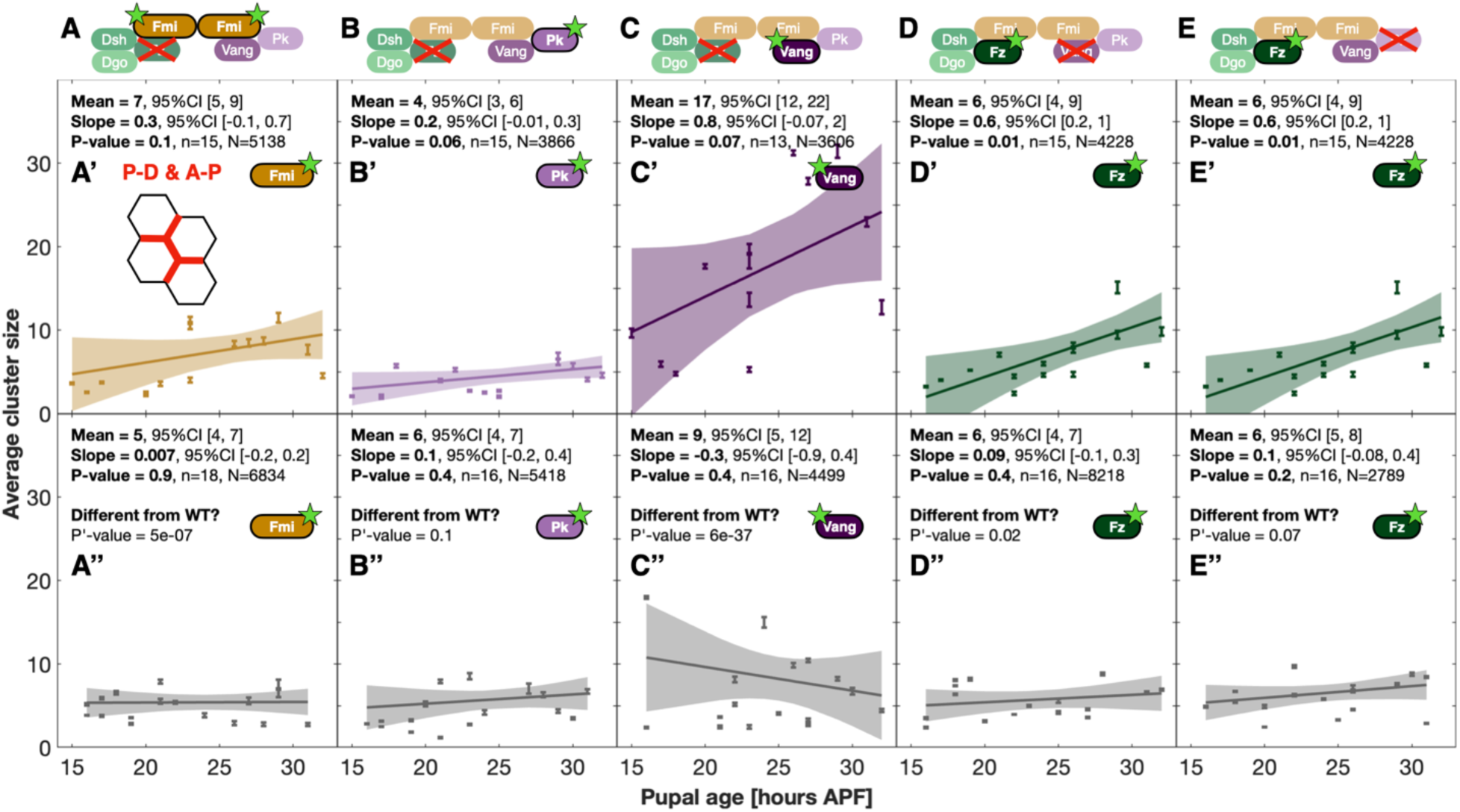
PCP clusters form in null mutants. (A-E) Schematic of the loss-of-function experiments. **(A’-E’’)** Cluster size distributions in wild-type (colored, **A’-E’**) compared to mutant conditions (gray, **A’’-E’’**), when examining all cell boundaries. **(A’-E’)** Wild type (the union of the data from Figures 2 and S2). **(A’)** Fmi::eGFP, **(B’)** Pk:eGFP, **(C’)** Vang::eGFP **(D’)** Fz::eGFP **(E’)** Identical to (D) **(A’’-E’’)** Loss-of-function data including all cell boundaries. **(A’’)** Fmi::eGFP in *fz^R52^/ fz^R52^*, **(B’’)** Pk::eGFP in *fz^R52^/ fz^R52^*, **(C’’)** Vang::eGFP in *fz^R52^/ fz^R52^*, **(D’’)** Fz::eGFP in *vang^stbm6^/ vang^stbm6^*, **(E’’)** Fz::eGFP in *pk^pk-sple13^/pk^pk-sple13^*. Differences in cluster sizes are most pronounced in wild-type late pupal ages, whereas little difference is found between wild-type and mutant conditions at earlier pupal ages. Dots, error bars, Mean, Slope, P-value, *n* and *N* as in Figure 2. P’ = significance of the difference between loss-of-function and corresponding wild type conditions by linear mixed-effects model test. For sample images of the loss-of-function conditions, see Figure S6.

Though we did not measure a failure to form clusters in the three null mutant condition examined, we cannot rule out that unmeasured proteins were not recruited to clusters for a given null mutant. As discussed in the companion study, the observations that cells lacking Fmi retain the ability to polarize, and that both Fz and Vang cluster in those cells, imply both that some level of cell-autonomous proximal and distal subcomplex formation occurs in the absence of Fmi and that these subcomplexes are competent to mediate feedback required for polarization^37^. Thus, clustering is robust to the absence of Fmi, Fz, Vang, and Pk, even though each is necessary for the growth of cluster sizes and tissue-level polarization.

### Growth of larger clusters is required for downstream signaling

Removal of Fz, Pk, or Vang does not eliminate clustering but instead inhibits both cluster growth and polarization. We therefore hypothesized that the assembly of larger clusters might be necessary to generate reliably polarized clusters and cells. However, an alternative explanation is that other functions of the missing proteins in the null mutants might account for the failure to polarize. We therefore sought a way to disrupt clustering as specifically as possible while leaving other functions intact.

To do so, we leveraged a well-defined structure/function relationship. The DIX domain of Dsh is known to oligomerize and is thought to mediate Dsh clustering *in vivo*^31,47,48^. A previous study characterized an allelic series of single amino acid substitutions in mouse Dishevelled2 (Dvl2) that disrupt oligomerization to varying degrees by disrupting antiparallel inter-strand interactions^31^. Based on these data, we created orthologous *Drosophila dsh^DIX^* point mutants expected to partially or entirely block Dsh oligomerization. Each, along with a wild-type control, was made on a transgene driven by the *Dsh* promoter and placed on a *dsh^v26^* null chromosome.

Three mutants, Dsh^N80D^, Dsh^N80A^, and Dsh^G63D^, produce a range of polarity phenotypic strengths, with Dsh^N80D^ being the weakest, Dsh^N80A^ being stronger, and Dsh^G63D^ slightly stronger still (Fig. 5A-D). We assess that Dsh^G63D^ is the strongest because, while hemizygous males of each were viable, homozygous females of Dsh^N80D^ and Dsh^N80A^ were viable, but Dsh^G63D^ females were not, and Dsh^G63D^ male wings consistently display notches consistent with Wingless (Wg) signaling defects not seen in the Dsh^N80D^ and Dsh^N80A^ wings, suggesting a progressively stronger effect on the Wg pathway as well. All three mutants express at levels that mediate sufficient Wg signaling to support some viability through pupal and adult stages. Dsh^N80D^, and Dsh^G63D^ express at levels similar to wild type, while Dsh^N80A^ expresses at a lower level (Fig. S7). Therefore, with the caveat of lower expression of Dsh^N80A^, these mutants form an allelic series with respect to PCP phenotype that mirrors the previously observed effects of their Dvl analogs in an *in vitro* Wnt signaling assay^31^.

**Figure 5:**
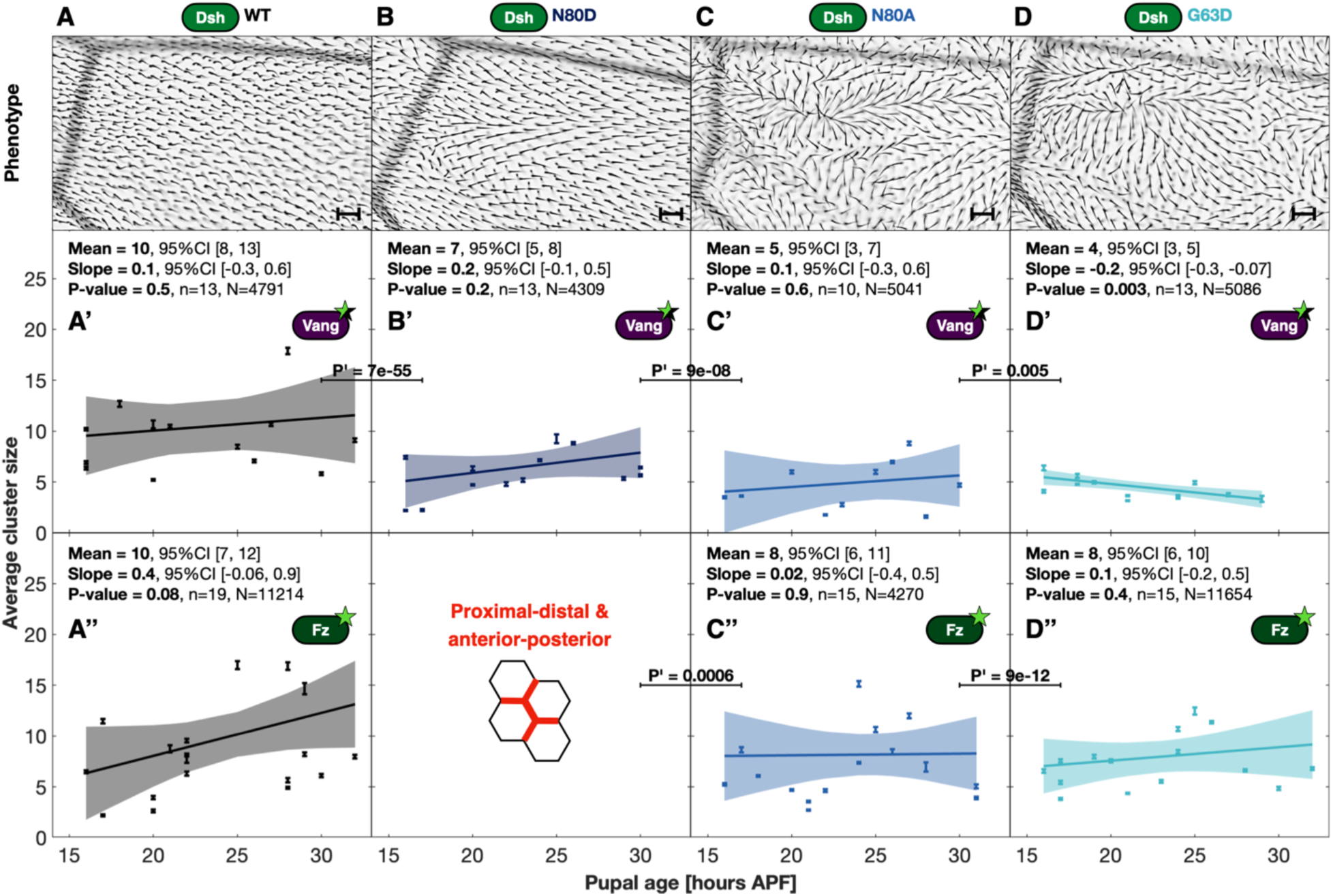
Growth of larger clusters is required for downstream polarity readout. (A-D) Trichome polarity patterns on the adult *Drosophila* wings near the posterior crossvein in hemizygous males expressing wild type or DIX mutant Dsh from a transgene on a *dsh^v26^* (null) chromosome: **(A)** Wild type (WT, control), **(B)** Dsh^N80D^, **(C)** Dsh^N80A^, **(D)** Dsh^G63D^. All scale bars are 20 μm. **(A’-D’)** The average number of heterozygous tagged Vang::eGFP monomers in clusters along the entire cell periphery in the **(A’)** WT (control), (B’) Dsh^N80D^ **(C’)** Dsh^N80A^, and (**D’)** Dsh^G63D^ conditions. **(A’’-D’’)** The average number of homozygously tagged Fz::eGFP monomers in **(A’’)** WT (control), **(C’’)** Dsh^N80A^, and (**D’’)** Dsh^G63D^ conditions. This experiment was not performed for Dsh^N80D^. Note the severity of the hair polarity phenotype correlates with reduced cluster sizes for Vang, and to a lesser extent, Fz. **(A’-D’’)**. Growth of Vang and Fz in WT **(A’, A’’)** might be expected, but because we are combining measures of proximal-distal where growth is significant (Figure 2) and anterior-posterior junctions, where it is not significant (Figure S2), significance is not reached here. Dots, error bars, Mean, Slope, P-value, *n* and *N* as in Figure 2. P’ = significance of the difference between two conditions by linear mixed-effects model test (C’’ compares to A’’).

To assess their impact on clustering, we introduced the Vang::eGFP probe into the three Dsh mutants. Viability limitations prevented us from creating flies with homozygous Vang::eGFP, so Vang::eGFP was instead used as a heterozygous probe, which allowed us to compare relative molecular counts. We observed that cluster formation still occurs in each mutant, but that Vang cluster size is reduced in each of the mutants relative to wild-type, with the extent of reduction roughly proportional to the strength of the polarity defect (Fig. 5A’-D’). Little or no cluster growth over time was observed, though our sensitivity for detecting growth with heterozygous probes was low, as little growth in the Dsh^WT^ control could be seen.

We also introduced Fz::eGFP as a homozygous probe and counted it in the presence of Dsh^N80A^ and Dsh^G63D^ (Fig. 5A’’-D’’). Again, Fz is incorporated into clusters, though of modestly decreased size, and no growth over time was observed in the Dsh^DIX^ mutants.

We conclude that Dsh^DIX^ mutants that selectively impair oligomerization while leaving other functions of the Dsh protein intact allow PCP clusters to form, but impair their ability to grow into larger clusters. These effects are proportional to their ability to mediate tissue-level polarization across alleles of varying strength. These data are consistent with the hypothesis that acquisition of large cluster sizes is required to effectively create cluster, cellular and tissue asymmetry.

### Two-color imaging reveals rough proportionality between pairs of components

Our analyses to this point determined the cluster size distributions of individual components. However, these assays could not reveal the degree to which the numbers of different components correlate within individual clusters. To do so, we created flies that uniformly express mScarlet::Vang plus Fmi::eGFP, Fz::mScarlet plus Vang::eGFP, and mScarlet::Vang plus Pk::eGFP (Fig. 6A-C). We then quantified the intensities of mScarlet and eGFP in each identifiable fluorescent punctum (Fig. 6A’-C’). (The properties of mScarlet did not permit reliable molecular counting.) Despite the expected noise due to the limitations of this quantification approach, for each combination, there is rough proportionality, indicating that, at least for Vang, Fmi, Fz and Pk, smaller clusters are smaller for each component, and larger clusters are larger for each component when assessing the summed P-D boundaries. The modest correlation we observe is consistent with a variable stoichiometry, as previously proposed^36^. The same relationship holds true for the A-P clusters (Fig. S8).

**Figure 6:**
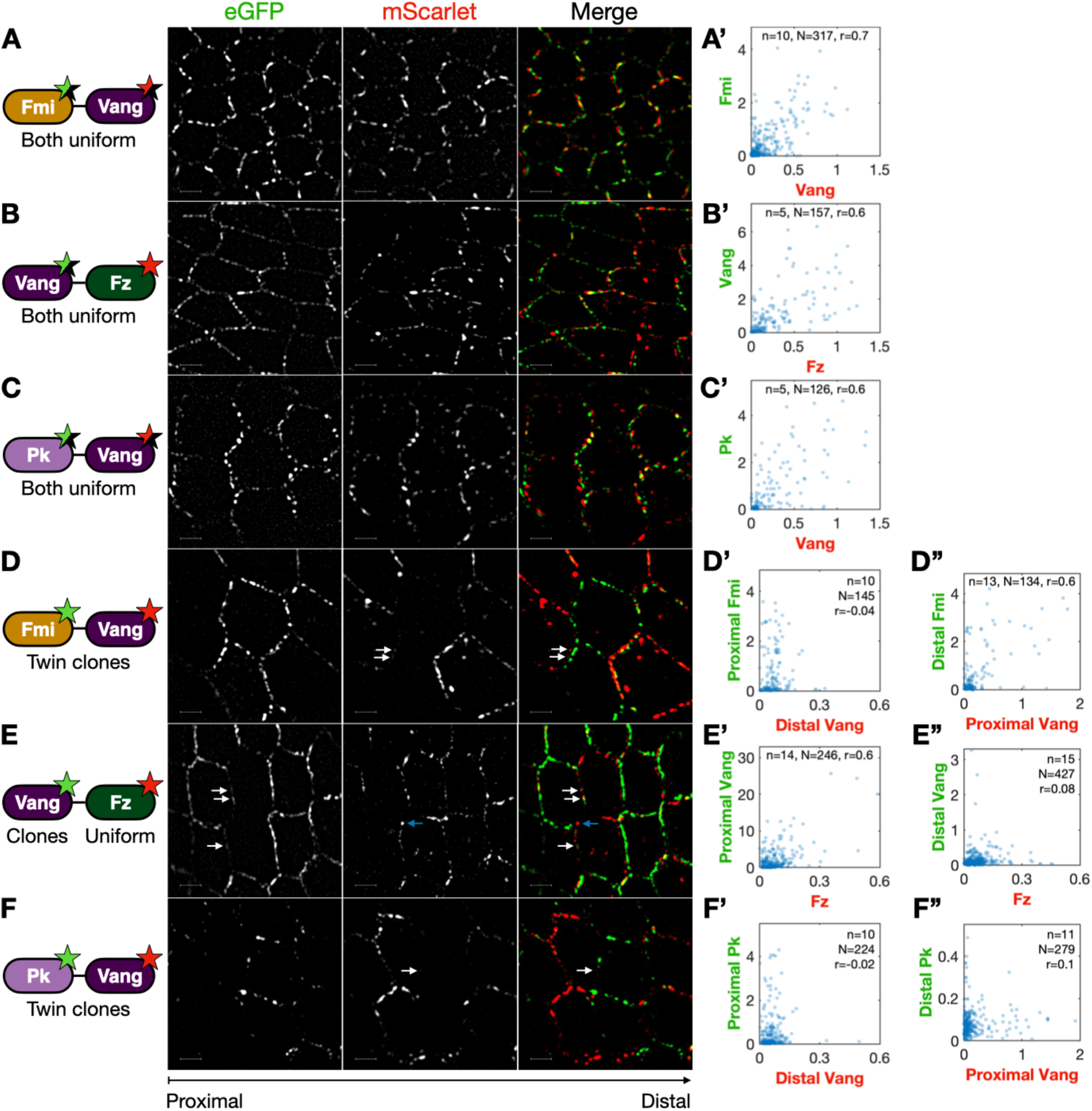
Two-color imaging reveals that large clusters are more asymmetrical and more likely to be oriented correctly. TIRF microscopy sample images of uniform expression of **(A)** heterozygous Fmi::eGFP, heterozygous mScarlet::Vang, 30 hr APF, **(B)** heterozygous Vang::eGFP, homozygous Fz::mScarlet, 24 hr APF, and **(C)** heterozygous Pk::eGFP, heterozygous mScarlet::Vang, 30 hr APF. **(A’-C’)** Analyzed intensities at paired subcomplexes along proximal-distal boundaries (details: see Methods). Each blue dot represents one (light blue) or more (dark blue) clusters. **(D)** Twin clones of Fmi::eGFP, mScarlet::Vang, 32 hr APF. **(E)** Clones of Vang::eGFP in a uniform Fz::mScarlet background, 30 hr APF. **(F)** Twin clones of Pk::eGFP, mScarlet::Vang, 29 hr APF. Analyzed intensities at paired subcomplexes along proximal or distal boundaries as labeled: **(D’)** Proximal Fmi vs. distal Vang. **(D’’)** Distal Fmi vs. proximal Vang. **(E’)** Proximal Vang vs. uniform Fz. **(E’’)** Distal Vang vs. uniform Fz. **(F’)** Proximal Pk vs. distal Vang. **(F’’)** Distal Pk vs. proximal Vang. *n* is the number of wings imaged. *N* is the total number of clusters analyzed. *r* is the correlation coefficient. All two-color wings imaged 27 hr ± 3 hr APF. White arrows point at examples of distal Vang. Blue arrow points to an example of strong Fz signal without detectable distal Vang. Note, the varying *x*- and *y*-scales, and the asymmetry degree of Fmi, Vang and Pk, matches the one found in Figure 3. All scale bars are 2 μm. For anterior-posterior boundaries, see Figure S8. We note that the method used to analyze these two-color images pairs subcomplexes with a detected signal in both channels within a defined distance, and does not quantify intensities inside clusters that do not have a detected partner in the other channel. If unmatched clusters were included, the reported anticorrelations should still hold true, and would further strengthen that trend.

### Mathematical model of polarized cluster formation

A model that captures all reported interactions between the core PCP proteins would be cumbersome, yet still incomplete given the likelihood that relevant protein-protein interactions and post-translational modifications are presently not known. We therefore sought to define the simplest possible model that could capture our and previous observations, and in so doing, define a minimal set of rules that may underlie the induction of polarity.

After exploring multiple possibilities, we focused on models in which clusters grow and shrink stochastically and independently. In the main text, we describe a model (Fig. 7A) that captures a minimal set of properties necessary for polarization to occur. A more molecularly detailed model described in the Supplemental Information captures additional, known molecular interactions. The base model incorporates four essential properties: ***i)*** the probability of cluster growth and shrinkage is (approximately) independent of cluster size, ***ii)*** Vang-Pk and Fz-Dsh-Dgo exhibit mutual inhibition intracellularly^23,27^, ***iii)*** Fmi-Fz preferentially binds intercellularly with Fmi-Vang^23,49^, and ***iv)*** trafficking of Fz and Dsh introduces a modest bias toward the distal size of individual cells^50–52^. To implement these properties, in a simple model, we combine the six PCP components into Fmi-Fz-Dsh-Dgo (denoted *F*) and Fmi-Vang-Pk (denoted *V*) subcomplexes.

**Figure 7:**
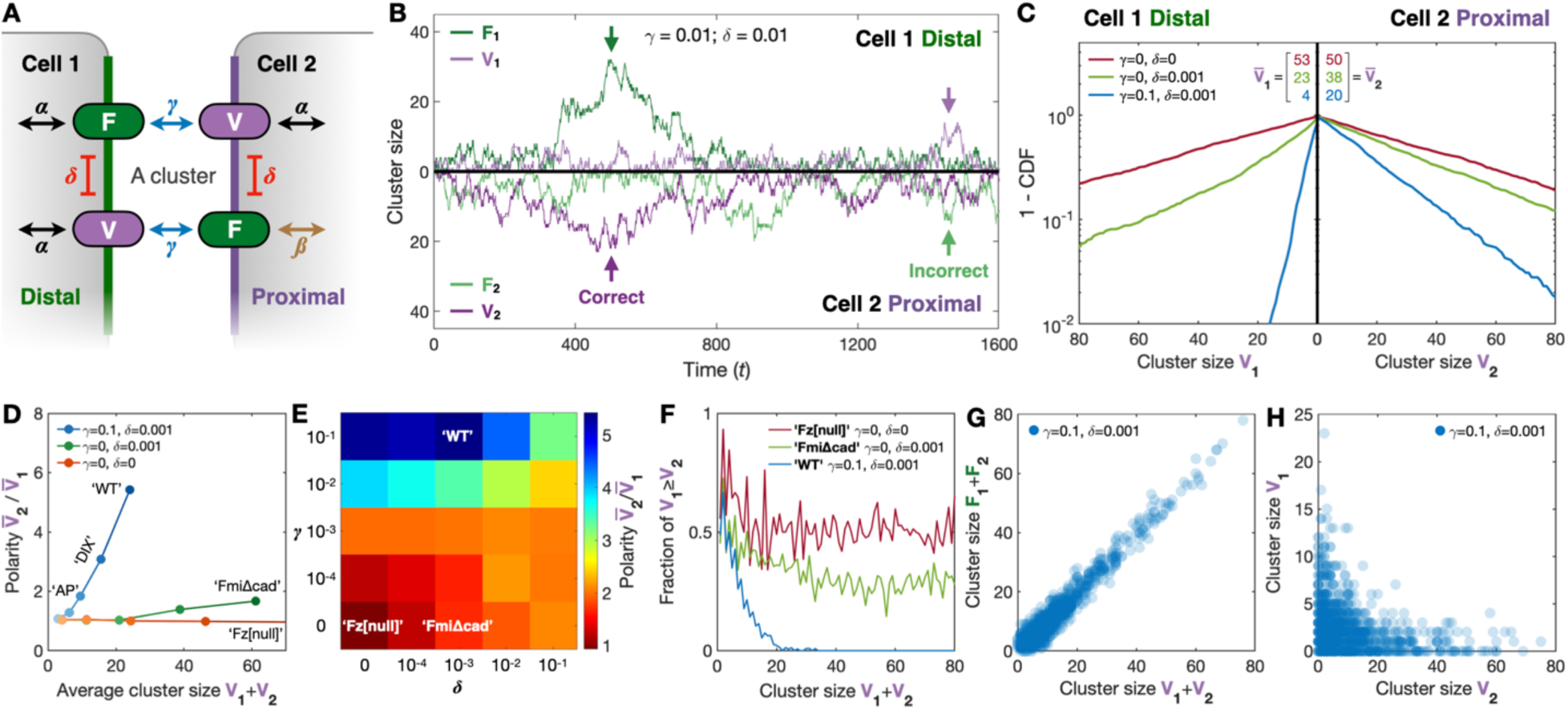
Mathematical model of planar cell polarity (PCP) cluster formation. **(A)** Schematic of a cluster with *F* and *V* subcomplexes that may accumulate on both sides of a junction based on the four parameters (*α*, *β*, *δ*, *γ*, see *Methods* for details). **(B)** Example simulation illustrating how the number of subcomplexes stochastically increases and decreases for a cluster on the two sides of the junction (indicated by the 0-line) as a function of time (updates). *F*_1_ indicates subcomplex *F* in cell 1, *F*_2_ subcomplex *F* in cell 2, *V*_1_ subcomplex *V* in cell 1, and *V*_2_ subcomplex *V* in cell 2. The arrows point at examples where the cluster is oriented correctly (dark colors) and incorrectly (light colors) relative to the defined tissue. **(C)** The distribution of *V*_1_ and *V*_2_ cluster sizes for *N* = 1000 clusters after *T* = 10^6^ updates for selected parameters. Blue mimics wild type with intercellular binding strength *γ* = 0.1 and intracellular inhibition strength *δ* = 0.001, green mimics FmiΔcad with *δ* = 0.001 and *γ* = 0, and red mimics loss-of-function e.g. fz[null] with *γ* = *δ* = 0. Note, the logarithmic *y* axis. CDF is the cumulative distribution function, and the overlines indicates the average number of subcomplexes for the three conditions. **(D)** The polarity as a function of the average cluster size by progressively increasing *α* (light-to-darker color gradient) for the same three conditions as in (C). Note how polarity scales with cluster size (blue line), how weak polarity can occur even when clusters are not communicating across junctions (green line), and how no polarity is achieved in null mutants (red line). **(E)** Phase diagram matching the right-most points in (E). **(F)** Fraction of incorrectly oriented clusters as a function of cluster size for the conditions in (C). **(G)** Uniform expression two-color simulations for the wild-type scenario; compare to Figure 6B’. Each dot represents one (light blue) or more (dark blue) clusters. **(H)** Clone boundary two-color simulation; compare to Figure 6F’ and F’’. Throughout the figure, the basal influx *α* = 0.99 (except for D, where *α* varies increases from 0.8 up to 0.99) and the biased influx *β* = 0.8. For (C-H), the number of simulated clusters *N* = 1000, the measurements occur at time *T* = 10^6^, For model variations and parameter scans, see the Appendix and Figures S9-S10, and for graphical simulations of the more complete model, see Movie S2.

These four features are captured with four corresponding parameters: ***i)*** Subcomplexes grow with rate *α’* per unit time and decay with a somewhat larger rate (*τ’*, Fig. 7A), meaning that stepwise disassembly is more probable than growth (*α* = *α’*/*τ’* < 1, Fig. 7B). This condition prevents runaway cluster growth and yields exponential distributions in cluster sizes (Fig. 7C)^40^. ***ii)*** The presence of *F* or *V* in the cluster inhibits the influx of the other subcomplex intracellularly (*δ*), i.e., the more *F* there is in the cluster, the less likely it becomes for *V* to enter that same side^26,27,53^. ***iii)*** Within a cluster, the subcomplexes *F* and *V* interact intercellularly (*γ*) such that the number of *V* and *F* subcomplexes tend to equalize on opposite sides of the cell-cell boundary, e.g., excess *F* on one side increases the probability of *F* to leave and the probability of *V* to enter on the opposite side in the next update, and vice versa for excess *V*. ***iv)*** We model an influx bias (*β*) that causes *F* to incorporate at a slightly smaller rate on the proximal side relative to the distal side, thereby breaking cluster symmetry.

Random, unordered incorporation and removal of cluster components (Fig. 7B) replicates the observations that cluster composition lacks fixed stoichiometry (^36^ and this study). We do not include self-promotion or -inhibition of *F* or *V* (i.e., the probability of adding or subtracting *F* or *V* is independent of the size of the cluster) since doing so would result in non-exponential cluster size distributions. The model in the main text does not capture the molecular interactions within the *F* or *V* subcomplexes, which can potentially be complex (see Appendix and Fig. S10).

The fraction of *V* or *F* on the incorrect side is primarily determined by *β* and *γ*, while the amount on the correct side, and therefore the overall size of clusters, is determined mainly by *α* (Fig. 7D). Setting *γ* and *8* to zero isolates *F* and *V* from each other, approximating Fz and Vang null mutants; this eliminates asymmetry while preserving the exponential distribution of the remaining component (Fig. 7C). If one assumes that Dsh^DIX^ mutants diminish the ability of components to oligomerize, one can approximate this condition by decreasing *α*; modestly reducing *α* causes cluster size to shrink and asymmetry to be reduced, as is seen experimentally (compare Fig. 7D to Fig. 5, see also Fig. S9). In contrast to the influx bias at P-D boundaries, A-P boundaries are not subject to this asymmetry and therefore remain on average symmetric. Setting *α = β* simulates this condition and results in symmetric clusters on the population level (Fig. 7D). Interestingly, as *δ* gets stronger, the size of complexes decreases, but if *δ* = 0, *F* will polarize but *V* will not polarize at all (Fig. S9). Therefore, it is likely that a moderate level of mutual inhibition occurs *in vivo*. In the companion manuscript, we show the surprising result that individual cells lacking Fmi, as well as cells expressing a truncated Fmi that cannot form *trans* dimers, still polarize. Modeling this condition by setting *γ* = 0 while keeping *δ* > 0 predicts that cells polarize, though less strongly than in wild-type (Fig. 7D and Fig. S9). A distillation of these results can be visualized in a phase plot (Fig. 7E).

Because the model posits that monomers of any core species can stochastically enter or leave a cluster on either side, the asymmetry of a particular core protein within any cluster can vary from perfectly symmetric to fully asymmetric. Experimentally, we indeed observe non-zero amounts of Fz, Pk, and, to a greater extent, Vang on the side where they are not dominant, consistent with such possible distributions. The model makes two important and non-intuitive predictions concerning cluster asymmetry in wild type (Fig. 3). First, the probability of correct cluster orientation increases with cluster size in a non-linear fashion (Fig. 7F), and second, large clusters are, on average, expected to be more asymmetric than smaller clusters (Fig. 7D). Increased fidelity in cluster polarization with cluster growth and decreased probability of reversed polarization would both be expected to produce a more coherent signal to downstream effectors.

While considerable insight can be derived from this simplified mathematical model, a more complex version can be used to describe the behavior of individual proteins rather than *V* and *F* subcomplexes, demonstrating the feasibility of stochastic unordered addition and subtraction of monomers and of the various interactions modeled by the equations, to reflect the underlying molecular biology (see Appendix, Fig. S10, and Movie S2).

### The degree of asymmetry and likelihood of correct cluster orientation increase with cluster size

To test the modeling predictions that the degree of asymmetry and probability of correct cluster orientation increase with cluster size, we returned to high resolution imaging. In so doing, we aimed to confirm that distal Vang indeed occurred within defined PCP clusters. If so, we reasoned that distal Vang could, in principle, indicate ***i)*** the presence of clusters with reversed orientation, ***ii)*** populations of distal and proximal Vang simultaneously in the same clusters, or both.

We first asked if the distal Vang clusters are indeed included in intercellular complexes or if they form independent of the other core proteins. To address this question, we analyzed interfaces between twin clones expressing mScarlet::Vang and Fmi::eGFP in adjacent cells (Fig. 6D), thereby unambiguously identifying signal on proximal and distal sides of the junction. Where distal Vang was present, it was almost always accompanied by proximal Fmi, suggesting that distal Vang is associated with Fmi homodimers spanning the intercellular junction.

Importantly, distal Vang was most often associated with small proximal Fmi clusters (Fig. 6D’). In contrast, per cluster, proximal Vang counts were roughly proportional to distal Fmi counts (Fig. 6D’’). Numerous large proximal Fmi clusters were not associated with distal Vang, implying that not all intercellular complexes contain distal Vang. These data indicate that distal Vang tends to exist within small clusters and tends not to be part of larger clusters.

We next explored whether distal Fz accompanies distal Vang. Imaging Vang::eGFP clones in a uniform Fz::mScarlet background (Fig. 6E), we saw that most distal Vang colocalized with Fz. Since almost all Fz was found to be on the distal side (Fig. 3B,B’,B’’), we infer that at least some distal Vang exists in clusters with distal Fz. Importantly, the amount of distal Vang anticorrelates with the amount of Fz (Fig. 6E’’), confirming that distal Vang tends not to be part of large clusters. As predicted, proximal Vang more closely correlates with Fz (Fig. 6E’). We similarly assayed the relationship between distal Vang and proximal Pk (Fig. 6F). At interfaces of twin clones expressing mScarlet::Vang and Pk::eGFP, we again see that the amount of distal Vang anticorrelates with the amount of proximal Pk (Fig. 6F’). These results support the presence of a few asymmetrical clusters oriented the wrong way, along with a somewhat larger number of correctly oriented clusters with a few molecules of distal Vang and larger numbers of distal Fz and proximal Vang. The largest Fz-containing clusters have very little, if any, distal Vang.

These results confirm key predictions from the mathematical modeling. If *F* and *V* are understood to represent full sub-complexes (Fmi-Fz-Dsh-Dgo and Fmi-Vang-Pk, respectively), then the correlations between proximal Vang and distal Fmi, and proximal Vang and Fz are as predicted (compare Fig. 6D’’,E’’, *r* ≥ 0.6, to Fig. 7G). The anticorrelations between distal Vang and proximal Fmi, distal Vang and Fz, distal Vang and proximal Pk, and proximal Vang and distal Pk (compare Fig. 6D’,E’’,F’’, r ≤ 0.1, to Fig. 7H) are likewise as predicted. We likewise know from confocal imaging that only adequately detects the larger clusters that most larger clusters are correctly oriented (see for examples refs. ^27,54^).

We interpret these results to indicate that a non-trivial fraction of small clusters may be incorrectly oriented, but during cluster growth, distal Vang and distal Pk tend to be excluded, and distal Fz and proximal Pk accumulate, such that the clusters that grow largest are the most likely to be both correctly oriented and highly asymmetric. Since the largest clusters are both most likely correctly oriented and least mobile (Fig. S4), we can also infer that correctly oriented clusters tend to be less mobile than their incorrectly oriented, smaller, counterparts. These relationships are expected to be stochastic, and it is likely true that some correctly polarized clusters containing larger amounts of distal Fz and proximal Pk (and, by inference, proximal Vang) may also contain some amount of distal Vang.

## DISCUSSION

Our measurements allow us to draw several detailed mechanistic conclusions: ***i)*** Exponential distributions of cluster sizes for all six core proteins are consistent with the simplest possible growth law, namely on- and off-rates that are independent of the cluster size; ***ii)*** clustering is observed in all null mutants examined, indicating that no single core protein is responsible for cluster formation; ***iii)*** a Dsh oligomerization mutant reveals that growth of larger clusters is required to achieve cell and tissue polarization. These findings led us to propose a simple model for PCP cluster assembly that incorporates each core protein stochastically. The model predicts, that ***iv)*** as clusters grow, they become increasingly polarized and increasingly likely to be correctly oriented; this is confirmed in our 2-color experiments showing that the previously unappreciated populations of distal Vang, distal Pk, and proximal Fz predominantly occur in small clusters, implying that clusters bearing misincorporated components are less likely to grow than those with correctly oriented components. We conclude that a simple set of rules governing interactions during assembly cause stochastic cluster assembly trajectories to lead from small, randomly oriented, clusters to large, highly asymmetric and correctly oriented clusters. The interactions we propose here allow the PCP pathway to produce a reliably polarized output despite a weak and noisy biasing input.

A complex web of protein-protein interactions and post-translational modifications likely contribute to PCP signaling. Here, we propose that these multifarious contributions converge on a limited set of rules that are sufficient for polarization (Fig. 7). Multiple potential models for cluster growth, such as fixed scaffolding, fission and fusion, and coarsening, are inconsistent with the observed exponential distributions^4,38^. Instead, single-exponential distributions imply that the probability of a monomer of any core protein entering or leaving a cluster is essentially independent of cluster size^40^. One possibility is that clusters behave as filaments to which monomers are added or subtracted only from one or both ends, as in actin and microtubule filaments. *In vivo* structural studies of PCP clusters will be required to test the filament hypothesis.

A prior study used fluorescence recovery after photobleaching (FRAP) to evaluate the non-exchanging, stable fraction of core proteins in “puncta” and “non-puncta” identified by confocal microscopy^28^. It is now evident that much of the “non-puncta” consists of smaller puncta, which was previously acknowledged as a possibility^28^. Regardless of this caveat, mechanisms involving stepwise, sequential addition and removal of individual proteins to PCP clusters, for example in a filament-type model, would predict that the non-exchanging fraction of proteins as measured by FRAP would increase as a function of cluster size—a result that is recapituated by our kinetic model.

We show that along the P-D polarization axis, whereas most Vang was known to localize proximally, a population of Vang molecules exist in distal clusters (Fig. 3D-D’). This observation, together with data in a companion manuscript^37^, suggests a potential mechanism contributing to symmetry breaking. In the companion manuscript^37^, it is shown that Vang associated with Fz has the propensity to recruit Pk, but that Pk recruitment is inhibited by the Fz-dependent recruitment of Dsh, which competes with Pk for binding to Vang^30,55,56^. A parsimonious explanation is that the presence of distal Vang reflects the stochastic dynamics of cluster assembly, and that any consequence of its presence is inhibited by Dsh blocking its ability to recruit Pk. Furthermore, distal Vang is less likely to be observed in larger clusters, suggesting that this competition also tends to limit Vang accumulation on the distal side. This competition likely serves as a symmetry breaking mechanism, and may be the mechanism underlying one part of negative feedback. This is discussed in more detail in Weiner et al.^37^

In this study, we tested the hypothesis that the growth of larger clusters is necessary for PCP signaling. This hypothesis harmonizes with our observation that the largest clusters along P-D boundaries are observed when apparent cellular asymmetry is at its greatest before it is read out at the time of pre-hair formation. To further probe the importance of cluster size, we used a series of mutations in the Dsh DIX oligomerization domain that tune the contribution of Dsh to cluster growth. These three mutants decreased cluster size proportionately to their polarity phenotypes (Fig. 5), providing evidence that larger average cluster sizes are required for PCP signaling. This did not answer the question, however, as to why larger clusters are necessary or how these clusters are read out.

Several non-exclusive models might explain how the formation of sufficiently large clusters is required for successful PCP signaling. Large clusters may form more effective and more stable scaffolds to signal downstream events. In this case, even abundant, well-polarized smaller clusters would be less effective in transducing a polarity signal. Alternatively, cluster growth may be associated with an increasing probability of strong asymmetry. Finally, clustering may be associated with increasing probability of correct orientation. The latter two relationships are predicted by our mathematical model, and confirmed by observation in our 2-color experiments.

To explore potential mechanisms that might capture our observations and those from prior studies, and to perhaps predict other features, we developed an abstracted mathematical model. The model provides a useful framework for inferring principles of cluster polarization while intentionally avoiding describing specific molecular interactions, about which too little is known to model in a meaningful way. The model, incorporating cell-autonomous mutual inhibition between proximal and distal complexes (*δ*), a tendency to equalize proximal and distal complexes in adjacent cells (*γ*), and an input bias to initiate symmetry breaking (*α* * *β*), successfully captures existing observations, providing evidence that these features are at least plausibly sufficient to describe polarization.

Furthermore, these results suggests that no fixed order of component addition to or subtraction from clusters need be prescribed, as our experimental results are replicated in model simulations that do not make such an assumption. Consistent with this notion, the assumption that Fmi is an obligatory scaffold for complex formation and symmetry-breaking is falsified by the observation in the companion manuscript of clusters of Vang and clusters of Fz in cells that polarize despite expressing no Fmi^37^.

The model, and our data, indicate that polarization occurs autonomously at each PCP cluster. Live-cell imaging indicates that the initiation and evolution of each cluster most often occurs independently, as we observe relatively few interactions between clusters on a time scale of an hour (Movie S1). Whether clusters are seeded at yet unidentified spatial landmarks or if they seed randomly remains to be determined.

Our model does not explicitly define time scales of on/off rates for *F* and *V* or individual core proteins, or of the life cycle of individual clusters. However, some observations place constraints on these values. Previous FRAP experiments suggest that the combined on/off rate for exchangable molecules occurs on a minutes timescale, which is somewhat slow compared to the typical rate of exchange of individual molecules, but is consistent with a stochastic addition and subtraction process proceeding through many steps. Our timelapse movies are expected to most readily visualize larger clusters, and the persistence of visible individual clusters in minutes to hours (Movie S1) is consistent with our simulations showing transient growth and decay (Fig. 7B); a life cycle on this timescale would allow polarity to remodel as cell junctions remodel. Our data do not allow us to distinguish between the possibilities that the growth in average cluster size between 15 and 32 hr APF reflects a steady state not yet reached or if it indicates a change in one or more parameters governing interactions (e.g. *α*) over that time frame.

The most salient result of our modeling was the prediction that as clusters grow, they become both more asymmetric and more likely to be correctly oriented. This prediction was validated through two-color imaging, and provides a compelling explanation for the necessity of clustering in polarization.

In summary, our data indicate that cluster formation plays a key role in PCP signaling, and that the emergent properties of a dynamical cluster assembly mechanism following simple rules can produce reliably polarized clusters. Conceptually, our study provides compelling evidence for the hypothesis that cluster growth provides a simple but highly effective mechanism of error correction that can amplify modest biases in Fz and Dsh trafficking into highly efficient polarization. We propose that cluster growth thus provides a powerful means of amplifying weak and noisy inputs into a robust cellular output, in this case cell and tissue-level polarization.

## METHODS

### Genotypes

Wild type uniform expression experiments

Figs. 1C, 2A, S1A, S2A, S3A: w; P[acman]-eGFP::Dgo, FRT40A, dgo^380^

Figs. 1D, 2B, S1B, S2B, S3B: yw, dsh^V26^, FRT18; P[acman]-eGFP::Dsh

Figs. 1E, 2C, 4D’, 4E’, S1C, S2C (dark colors), S3C: w; +/+; Fz::eGFP

Figs. 1F, 2D, 4A’, S1D, S2D (dark colors), S3D: w; Fmi::eGFP

Figs. 1G, 2E, 4B’, S1E, S2E (dark colors), S3E: w; eGFP::Pk

Figs. 1B, 1H, 2F, 4C’, S1F, S2F (dark colors), S3F, Movie S1: w; P[acman]-Stbm::eGFP, FRT40A, Vang^Stbm6^ (P[acman]-Stbm::eGFP, a.k.a P[acman]-Vang::eGFP; ref ^36^)

Wild type clone boundary experiments

Figs. 3A-A’’, S2C (light colors), S4A-A’, S5A-A’: UbxFLP; FRT42D Stan::eGFP / FRT42 arm-lacZ

Figs. 3B-B’’, S2D (light colors), S4B-B’, S5B-B’: UbxFLP; arm-lacZ FRT73,80 / Fz::eGFP FRT80B

Figs. 3C-C’’, S2E (light colors), S4C-C’, S5C-C’: UbxFLP; FRT42D eGFP::Pk / FRT42D arm-lacZ

Figs. 3D-D’’, S2F (light colors), S4D-D’, S5D-D’: UbxFLP; P[acman]-Stbm, FRT40A, Vang^Stbm6^ / P[acman]-Stbm::eGFP-LoxP, arm-lacZ, FRT40A, Vang^Stbm6^

Loss-of-function uniformly labeled experiments

Figs. 4A,A’’, S6A: w; Fmi::eGFP; Fz^R52^

Figs. 4B,B’’, S6B: FRT42D, pk::eGFP; Fz^R52^

Figs. 4C,C’’, S6C: P[acman]-Stbm::eGFP, FRT40A Vang^Stbm6^; Fz^R52^

Figs. 4D,D’’, S6D: Vang^Stbm6^; Fz::eGFP

Figs. 4E,E’’, S6E: FRT42D, pk^pk-sple13^; Fz::eGFP

Dsh[DIX] uniformly labeled experiments

Fig. 5A: Dsh-v5-6xmyc, dsh^V26,f36a^

Fig. 5B: Dsh^N80D^-v5-6xmyc, dsh^V26^

Fig. 5C: Dsh^N80A^-v5-6xmyc, dsh^V26^

Fig. 5D: Dsh^G63D^-v5-6xmyc, dsh^V26^

Fig. 5A’: Dsh-v5-6xmyc, dsh^V26,f36a^; + / P[acman]-Stbm::eGFP, FRT40A, Vang^Stbm6^

Fig. 5B’: Dsh^N80D^-v5-6xmyc, dsh^V26^; + / P[acman]-Stbm::eGFP, FRT40A, Vang^Stbm6^

Fig. 5C’: Dsh^N80A^-v5-6xmyc, dsh^V26^; + / P[acman]-Stbm::eGFP, FRT40A, Vang^Stbm6^

Fig. 5D’: Dsh^G63D^-v5-6xmyc, dsh^V26^; + / P[acman]-Stbm::eGFP, FRT40A, Vang^Stbm6^

Fig. 5A’’: Dsh-v5-6xmyc, dsh^V26,f36a^; Fz::eGFP

Fig. 5C’’: Dsh^N80A^-v5-6xmyc, dsh^V26^; Fz::eGFP

Fig. 5D’’: Dsh^G63D^-v5-6xmyc, dsh^V26^; Fz::eGFP

2-color experiments

Fig. 6A-A’, S8A: hsFLP; FRT42D,fmi::eGFP / FRT42D,mScarlet::Vang, Vang^Stbm6^ (no heat-shock)

Fig. 6B-B’, S8B: hsFLP; P[acman]-Stbm,FRT40A,Vang^Stbm6^ / P[acman]-Stbm::eGFP,FRT40A, Vang^Stbm6^; Fz::mScarlet (no heat-shock)

Fig. 6C-C’, S8C: hsFLP; FRT42D,pk::eGFP / FRT42D,mScarlet::Vang, Vang^Stbm6^ (no heat-shock)

Fig. 6D-D’’: hsFLP; FRT42D,fmi::eGFP / FRT42D,mScarlet::Vang, Vang^Stbm6^ (12 min 37 °C heat-shock at third instar larvae stage)

Fig. 6E-E’’: hsFLP; P[acman]-Stbm,FRT40A,Vang^Stbm6^ / P[acman]-Stbm::eGFP,FRT40A, Vang^Stbm6^; Fz::mScarlet (12 min 37 °C heat-shock at third instar larvae stage)

Fig. 6F-F’’: hsFLP; FRT42D,pk::eGFP / FRT42D,mScarlet::Vang, Vang^Stbm6^ (12 min 37 °C heat-shock at third instar larvae stage)

### Molecular counting

To determine how PCP clusters break symmetry, we developed a quantitative method to count the number of each of the six core proteins in individual PCP clusters. By analyzing bleaching traces obtained with a high frame rate in live *Drosophila* wing cells imaged using total internal reflection microscopy (TIRF), we quantified the intensity of single fluorophores in each punctum and calculated the molecular number in the corresponding PCP cluster.

### Sample preparation

Fly stocks were maintained on standard fly food. White pupae were selected at 0-3h after puparium formation (APF) and maintained at 25 °C until 15-32 hr APF. From each pupa, one wing sac was exposed by making a window in the pupal case^57,58^. A droplet of Halocarbon oil 700 (Sigma Life Science, refractive index 1.4, CAS No.: 9002-83-9) was placed on top of the exposed wing sac, and the wing was imaged at selected ages through a cover glass of thickness 1.5 (VWR Micro Cover Glass, selected 1 ounce, 18x18 mm, No. 1.5, VWR Cat No 48366-205).

### Image acquisition

Live wings were imaged on an inverted total internal reflection fluorescence (TIRF) microscope (Nikon TiE) with an Apo TIRF 100x oil objective lens and numerical aperture 1.49 (see also refs.^59,60^). Distal eGFP expressing wing cells were located using low intensity (1%) 473-nm OBIS laser (Coherent). For 1-color experiments (eGFP), the exposed wing cells were fully bleached at high intensity (50%) laser power using the 473-nm OBIS laser and a sequential exposure of 50 ms. For 2-color experiments (mScarlet + eGFP), the exposed wing cells were first fully bleached using a 532-nm Crustalaser, and then fully bleached using the 473-nm OBIS laser. Both channels were acquired with 50 ms sequential exposure. An emission filter at 514/30nm (Semrock Inc.) was used for eGFP, and a 593/40nm filter (Semrock Inc.) was used for mScarlet. The images were recorded on a Hamamatsu Orca Flash 4.0 camera (fixed pixel size of 64 nm), and the scope was controlled using Micromanager^61^ with Z-autofocus. The field of view varies from image to image due to variations in the wing area exposed. The number of frames acquired also varies from image to image due to variations in bleaching time needed, typically between 1,000 and 5,000 frames.

### Image analysis

One image was selected per wing imaged. This image was selected based on the maximum number of cells in focus and *xy* drift less than 2px during image acquisition. No further drift correction was applied. These initial evaluation steps were performed in ImageJ^62^. All other post-processing steps were performed using custom-built MATLAB scripts. Cell boundaries, based on manually drawn nine px-wide masks were manually grouped into A-P and P-D boundaries for uniform expression wild type experiments and D, P, A or P for clone boundary wild type experiments. For mutant wing cells, all boundaries were assigned the same category. In the case of overlapping masks, P-D boundaries take precedence over A-P boundaries. For 2-color imaging, multi-modal image registration was performed based on the intensities in the first frame of each channel and applied on all frames of the mScarlet channel. Image transformation of the mScarlet channel was performed by integer *xy* pixel-translation only. No rotation, re-scaling, or shear were applied.

### Locating clusters

For each image, the following steps were applied on all frames: to flatten the illumination and apply bleaching correction, a 2D Gaussian fit (2x2 pixels wide) was subtracted. To reduce camera pixel noise, the average intensity value within a moving box (3x3 pixels wide) was calculated. To minimize temporal noise, a directional moving median filter was calculated (with a maximum of 20 frames before and 0 frames after the current position). Sample images in Figs. 1, 3, and S6 are shown according to this protocol. To locate cluster centers inside the masked boundaries, local (5x5 pixels wide) maxima were identified. These pixel centers correspond to cluster centers.

Although we limit our counts to fluorophores inside manually masked membrane junctions (of width roughly 250 nm on either side of the junctions) and in a common *z*-plane, the method does not distinguish between monomers integrated into a cluster vs monomers that are not. We assume that highly mobile and therefore freely diffusing fluorophores account for a small fraction of identified counts, and that most fluorophores located in this way are incorporated into clusters. This places some constraints on the accuracy of our measurements that apply to both low density junctions (possibly overcounting monomers) and high-density junctions (possibly undercounting monomers hidden close to large bright clusters). Together, we expect these inaccuracies to minimally impact the final counts.

### 2-color intensities

To pair clusters in two different channels, the following approach was used on both channels: Inspired by the SRRF method^63^, spatial resolution was increased by subdividing each pixel into 2x2 pixels in the *xy* direction and assigning the acquired value to the 4 subdivided pixels (new pixel size = 32x32 nm). Like the approach listed above, the illumination was flattened by subtracting a 2D Gaussian fit everywhere (5x5 pixels wide). Spatial noise was minimized by calculating the mean intensity value within a moving box (also 5x5 pixels wide). Temporal noise was reduced by determining the accumulated intensity inside each pixel during the first 5 frames. Cluster centers were located inside both channels independently by identifying local maxima (5x5 pixels). The processed intensity inside the center pixel of each cluster was dilated (4x4 pixels). Any uniquely overlapping dilated pixels across the two channels are said to correspond to a cluster containing the two labeled proteins.

### Measuring cluster size

The following steps were applied for each cluster’s pixel center: a histogram of the pair-wise difference is calculated for all frames to all future frames. The power spectrum is applied to this histogram, and the highest peak in the intensity interval between 40 and 110 is found. As internal validation, single fluorophore blinking events were found to occur within this interval.

The highest peak indicates the most likely fluorophore intensity step size for that cluster. To estimate the number of fluorophores present in the first frame, the intensity of the first frame is divided by the local fluorophore step size and rounded down to the nearest integer. This counting method is inspired by refs. ^64–66^.

MATLAB image analysis scripts deposited at https://github.com/SilasBoyeNissen/Cluster-Assembly-Dynamics-Drive-Fidelity-of-Planar-Cell-Polarity-Polarization.

### Validation by pentamer counting

To validate our molecular counting method, we synthesized and counted a purified pentameric GFP protein according to the following protocol.

Cloning: A fragment with DNA corresponding to the Lyn kinase myristoylation site, monomeric enhanced GFP, and the cartilage oligomeric matrix protein coiled-coil domain^67^ separated with 3x GGS linkers with a 5’ NdeI and 3’ XhoI site and standard adaptors was synthesized by Twist Biosciences. The fragment was digested with NdeI and XhoI and ligated into the NdeI and XhoI sites in pET15b, 3’ of the His6-thrombin sequence. The thrombin site was subsequently changed to a TEV site via Gibson assembly-mediated mutagenesis using the following primers: MY0099 thrombin to tev rvs: 5’tccTCCTTGgaaATACAGGTTTTCgtgatgatgatgatgatggctgctg 3’ MY0100 thrombin to tev fwd: 5’GAAAACCTGTATttcCAAGGAggaggctgtataaaatctaaacgaaaagatggcg 3’

Expression and purification: Chemically competent BL21(DE3) E. coli were transformed with pET15b-His6-TEV-Lyn-GFP-COMP and plated onto LB agar with 100ug/ml ampicillin. A single colony was used to inoculate 20ml of LB with 100ug/ml ampicillin and cultured overnight at 37C. The entire 20ml preculture was added to 2L of LB with 100ug/ml ampicillin and cultured at 37C, shaking at 225rpm until OD∼1.5. Temperature was dropped to 18C and shaking dropped to 175rpm and culture was induced with IPTG added to a final concentration of 0.1mM. Culture was harvested after 18h of induction via centrifugation at 8000xg and pellets were snap frozen with liquid nitrogen. Pellets were thawed into 250ml lysis buffer (50mM Tris pH 7.5, 150mM NaCl, 30mM imidazole, 1mg/100ml DNAseI, 4mM MgCl2, 2.5mM beta-mercapto-ethanol, 150nM aprotinin, 1uM E-64, 1uM leupeptin) and lysed using an Avestin Emulsiflex C3. Lysate was clarified via centrifugation at 30,000xg for 30 minutes. Lysate was loaded via gravity over 4ml Cytiva Ni Sepharose Fast Flow resin pre-equilibrated with 50ml lysis buffer. Column was washed with 100ml wash buffer (20mM Tris pH 7.5, 300mM NaCl, 30mM imidazole, 2.5mM beta-mercapto-ethanol). Protein was eluted with elution buffer (20mM Tris pH 7.5, 1M imidazole, 150mM NaCl, 2.5mM beta-mercapto-ethanol). Protein was diluted 1:10 into storage buffer (20mM Tris pH 7.5, 150mM NaCl, 1mM DTT) and loaded over a MonoQ column, eluting with a gradient of high salt buffer (20mM Tris pH 7.5, 1M NaCl, 1mM DTT). Fractions were analyzed via SDS PAGE and subjected to additional purification via size-exclusion chromatography over a Superdex 200 column in storage buffer. Fractions were analyzed via SDS PAGE, pooled, frozen in liquid nitrogen, and stored at −80C.

Multiangle light scattering analysis: The molecular weights of the purified Lyn-GFP-COMP pentamers were determined by size exclusion chromatography coupled with inline multi-angle light scattering(MALS) using a Superdex 200 10/300 GL column attached to a UV detector followed by a DAWN Heleos-II and an Optilab T-rEX refractive index detector (Wyatt Technology, Santa Barbara CA). 100ul of purified protein at 2mg/ml was injected over the system equilibrated in 20mM Tris pH 7.5, 150mM NaCl, 1mM EDTA, 0.02% NaN3 at 25C. Calibration of the detectors was performed by measuring the signal of monomeric bovine serum at 70kDa. The absolute mass was determined over the course of the run using ASTRA 6 software (Wyatt Technology) using the signals from the MALS and the RI detectors, confirming the expected molecular mass for a pentamer (Fig. S11).

### Imaging adult wings

For Figure 5A-D, adult flies were washed in 70% ethanol overnight at 5 °C. Wings were dissected, mounted in DPX solution (Sigma Life Science, 06522-100 ml, lot BCBT5387), and imaged on a Nikon model Eclipse Ni-e (921092) with SPOT Software (v. 5.6).

### Statistical tests

For each wing, the average cluster size and its uncertainty are found by performing a non-linear regression model (a single exponential fit) on the experimentally found cluster size distribution. To assess whether the cluster size follows an exponential distribution, we used histograms and Q-Q plots for a visual inspection. The histogram of Vang::eGFP cluster size in the age interval 15-23 hr APF, for example, shows that the data (blue bars) closely follows the exponential density plot (red curve), indicating a good fit to an exponential distribution (Fig. S11A; data from Fig. 1). Similarly, the Q-Q plot for Dgo::eGFP cluster size in the age interval 24-32 hr APF demonstrates that the data points align well with the reference diagonal line, suggesting that the data matches the theoretical quantiles of an exponential distribution (Fig. S11B; data from Fig. 1). Together, these plots showed us that it is reasonable to assume the population distribution of cluster sizes for each wing follows an exponential distribution.

To test whether the average cluster size grows with age, a weighted linear regression model is performed on all wings with the same genotype. The mean value reported is now the predicted response of the linear regression model at 23.5 hr APF. The P-value reported is the t-statistic of the two-sided hypothesis test on the slope of the linear model. A P-value below 0.05 and a positive slope within error suggest that cluster size likely increases with age.

A linear mixed-effects model is used to test whether two genotypes have statistically different cluster sizes from each other. The model formula, Cluster size ∼ age + genotype + age * genotype, evaluates the effects of age and genotype on cluster size, including their interaction. We report the P’-value for the genotype term. A P’-value below 0.05 indicates that the cluster sizes likely depend on the genotype.

### pCas4-dsh-V5-6myc

The V5-6myc fragment, flanked by Xba1 sites, was generated by touch down PCR using pUAST-Rab35-6myc (Addgene #53503) as template.

Forward primer: TCTAGAAGCGGCACAGGCTCTGGCGGCAAACCGATTCCGAACCCGCTGCTGGGCCTGGATAGCACCAGTGGTGGATCCACCATGGAGCAAAAGCTC

Reverse primer: TCTAGAATACCGGTGATTACAAGTCCTCTTC

pCas4-dsh from ref. ^68^ was digested with Xba1 and the V5-6xmyc fragment appended. The resulting plasmid was transformed into *w1118* flies to generate random genomic insertions, and an X chromosome insertion was recombined onto a *dshv26 f36a* chromosome.

### DIX Domain Mutants

The pCas4-dsh-V5-6xmyc plasmid was mutated using TOPO Cloning to introduce conserved base changes analogous to those in Kan et. al.^31^, resulting in N80D, N80A, and G63D mutations to the DIX domain. These three plasmids were then transformed into *w1118* flies to generate random genomic insertions, and an X chromosome insertion was recombined onto a *dshv26 f36a* chromosome. Parentheses indicate the analogous mouse mutation from Kan et al. (2020)^31^.

A Zero Blunt TOPO Invitrogen kit introduced the following point mutations within the original pCasper4-dsh-V5-6myc plasmid.

G63D (G65D) F GCCGATTTCGATGTGGTCAAA. (GGT>GAT)

R GTCCATTGACTTGAAGAAGTAC

N80D (N82D) F GCCCTGCTTCGATGGGCGAGT. (AAT>GAT)

R AGTATGGTGGAGTCGTCGG

N80A (N82D) F GCCCTGCTTCGCTGGGCGAGT. (AAT>GCT)

R AGTATGGTGGAGTCGTCGG

PCR template pCasper4-dsh-V5-6myc (wild type control) with primers

F GGCAGCGCTGGCAGTGTGACC.

R TCAGCACATCGCTGCTACTCG.

(Primers are outside Mlu1 and Xho1 digest sites.)

Individual Primer sets were used to introduce DIX point mutations via PCR into the original pCasper4-dsh-V5-6myc template, and these point mutations were inserted into the TOPO vector between the EcoR1 sites. Both the TOPO product (Dsh containing DIX mutation) and pCasper4-dsh-V5-6myc plasmid were digested using the Mlu1 and Xho restriction enzymes and the resulting fragments were then ligated together, resulting in pCasper4-dsh^DIX^-V5-6xmyc. The prepared plasmid was sent to BestGene for injection. Random X insertions were recombined onto the *dsh^V26^* chromosome.

### Western blot analysis

Third instar female *Drosophila* larval wing discs were dissected in cold PBS and transferred to Eppendorf tubes containing cold PBS with protease inhibitors. The samples were centrifuged at 14,000 rpm for 4 minutes at 4°C, and the supernatant was discarded. The tubes were snap frozen in liquid nitrogen and stored at −80°C. For protein extraction, samples were thawed, and RIPA buffer was added. Wing discs were hand-homogenized with a pestle, followed by centrifugation at 14,000 rpm for 10 minutes. The supernatant was collected, and 4x loading buffer was added before boiling the samples for 5 minutes.

Twenty microliters of each sample was loaded onto a 10% SDS-PAGE gel, and proteins were separated by electrophoresis. The proteins were then transferred to a PVDF membrane at 100 V for 90 minutes in a cold room. The membrane was cut horizontally at the 50 kDa marker band, resulting in two sections. Both sections were blocked for 30 minutes in 5% BSA-TBSTw.

The upper section was incubated with a 1:1,000 dilution of monoclonal V5-Tag antibody in 3% BSA-TBSTw, while the lower section was incubated with a 1:20,000 dilution of monoclonal anti-β-actin antibody in 3% BSA-TBSTw. Membranes were incubated overnight at 4°C.

Following incubation, membranes were washed three times with TBSTw for 10 minutes each. The membranes were then incubated with HRP-conjugated secondary antibodies in 3% BSA-TBSTw for 30 minutes at room temperature, with a dilution of 1:10,000 for the V5-Tag antibody and 1:30,000 for anti-β-actin antibody. After five washes with TBSTw (5 minutes each), membranes were treated with enhanced chemiluminescence (ECL) solution for 2 minutes, drained, and imaged using the Bio-Rad ChemiDoc MP Imaging System.

### Generation of mScarlet::Vang flies

We constructed the pBDP-mScarlet::Vang plasmid using Gibson assembly, with the NEB HiFi assembly kit. We used pBDP (https://www.addgene.org/17566/) as backbone. Vang 5’, coding sequence (CDS), and 3’ sequences were cloned from the BAC attB-P[acman]-AmpR-mApple-loxP-stbm (ref. ^33^) mScarlet was obtained from pF3BGX-mScarlet (https://www.addgene.org/138391/). We integrated the mScarlet fragment after the Vang promoter region, right before the Vang CDS starting codon, followed by the linker sequence SGTGSG. We created the final plasmid using the following 5 PCR fragments:

Fragment 1: pBDP backbone. Primers: pBDP Fwd + pBDP Rev

Fragment 2: Vang 5’. Primers Vang5’ Fwd + Vang5’ Rev

Fragment 3: Linker-mScarlet. Primers Linker-mScarlet Fwd + Linker-mScarlet Rev

Fragment 5: Vang 3’. Primers Vang3’ Fwd + Vang3’ Rev

Primer sequences:

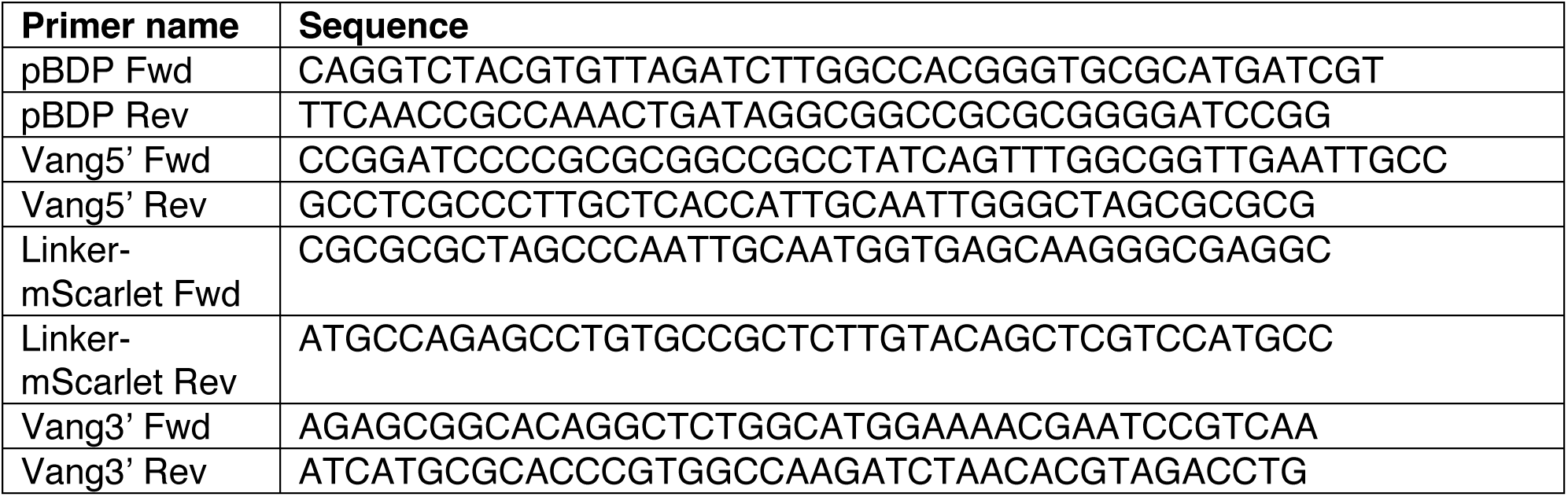

We inserted the plasmid using phiC31 into the attP landing site ZH-51C (BDSC #24482) and recombined it onto a *Vang^stbm6^* (null) chromosome.

### Math model

We introduce a Monte Carlo model consisting of two subcomplexes, *F* and *V*, that can oligomerize on both sides, 1 and 2, of a cluster spanning a cell-cell junction (Fig. 7A). Thus, as an example, *F*_2_ is the number of *F* subcomplexes that the cluster contain in cell 2. The evolution, *t*’, of the number of the two subcomplexes in the cluster is described by the rate equations:

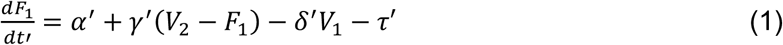

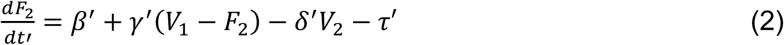

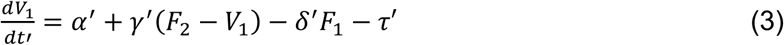

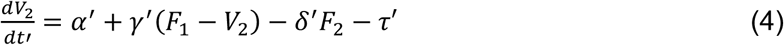

where *α’* is the on rate of *F*_1_, *V*_1_, *V*_2_, *β’* is the on rate of *F*_2_, *γ’* is the intercellular coupling strength, *δ’* is the intracellular inhibition strength, and *τ*’ is the rate at which the subcomplexes leave the cluster. These rate constants are unknown, and we assume they do not change with time nor are dependent on the state of the cluster.

We rescale the parameters by setting *t=t’ τ*’, *α=α’/τ*’, *β=β’/τ*’, *γ=γ’/τ*’, and *δ=δ’/τ*’, allowing us to reduce the rate equations to:

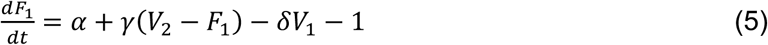

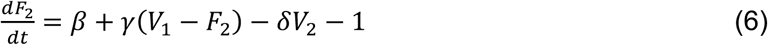

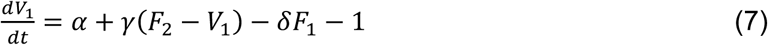

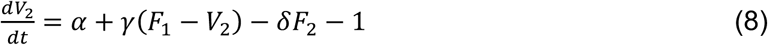

*β* breaks the symmetry of the cluster when *β* ≠ *α*. Thereby, *F* can be considered an input subcomplex that eventually breaks the symmetry of the output subcomplex. To avoid infinite cluster growth, the off-rate (*τ’*) is larger than the two on-rates (*α’* and *β’).* This condition implies that the number of monomers in the cluster performs a biased random walk towards 0 – the cluster is always about to collapse, and progressively larger clusters become exponentially less likely to occur. The parameter *γ* is the reduced intercellular coupling rate between different subcomplexes across the junction. A value of *γ* = 1 implies a 1:1 stoichiometry between the subcomplexes, whereas *γ* = 0 implies that the subcomplexes are uncoupled and uncorrelated. The parameter *δ* is the intracellular inhibition rate, which here increases linearly with the number of opposing subcomplexes within the cluster. For *δ* = 0, the growth of one subcomplex is not inhibited by the opposing subcomplex, whereas for *δ* = 1, the inhibition is maximal.

Initially, for time *t* = 0, the cluster does not contain any subcomplexes, meaning *F*_1_ = *V*_2_ = *F*_2_ = *V*_1_ = 0. We disallow subcomplexes of negative numbers to occur, by setting the resulting decay rate to zero when none are present. Otherwise, each following Gillespie model update stochastically selects a component to add or subtract based on the state of the cluster at that time. Thus, this model is a Markov process with no memory. The model runs as long as *t*<*T* and is repeated for *N* independent clusters. The model is implemented in MATLAB, and the source code has been deposited at https://github.com/SilasBoyeNissen/Cluster-Assembly-Dynamics-Drive-Fidelity-of-Planar-Cell-Polarity-Polarization.

## Acknowledgments

We thank members of the Axelrod lab for fruitful discussions. Thanks to David Strutt for providing fly lines and the Stanford Research Computing Center for providing computational resources. We thank Claire Tomlin and James Ferrell for fruitful discussions on the mathematical model and William Weis for advice in choosing the point mutations in Dishevelled. Thanks to Alexandre Ravel for his support in the initial attempts to image the Dishevelled mutations and to Gemma Landa for her support to image the counting control. Stocks obtained from the Bloomington Drosophila Stock Center (NIH P40OD018537) were used in this study.

This work was funded by NIH R35GM131914 (JDA), R35GM130332 (ARD), and NIH R01HL16929901 (JDA and ARD). SBN was supported by the Novo Nordisk Foundation (grant awards NNF20OC0059462 and NNF21CC0073729) and the Stanford Bio-X Program. MY was supported by the National Science Foundation Graduate Research Fellowship Program (grant no. DGE-1656518), and the Stanford Bio-X Bowes Fellowship. The funders had no role in study design, data collection and analysis, decision to publish, or preparation of the manuscript.

## Appendix

### A more complete math model

To describe the stochastic behavior of an entire core complex, we expand on the simplified model described in *Methods* to include five core proteins (Dsh, Fz, Fmi, Vang, and Pk). Along with the model schematic in Fig. S10A, the following 10 equations describe the rescaled rate equations of each component. Note that the two sides are symmetric except for Fz.

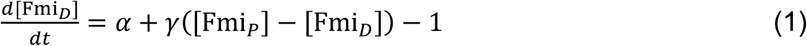

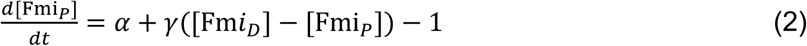

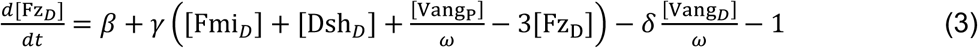

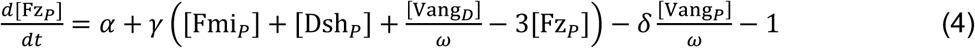

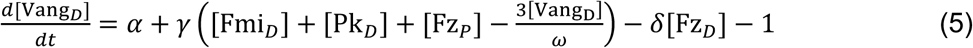

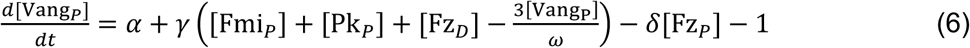

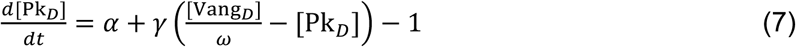

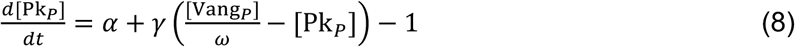

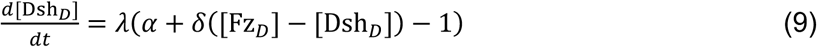

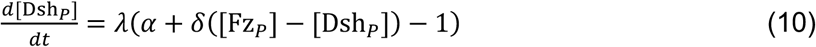

Like the simple model in the main text, *α* is the on-rate of proteins to a cluster. Here, it gives the average number of Fmi monomers in a cluster, and we anticipate that *α* grows with age, given the results in Fig. 2. *β* < *α* eventually breaks the symmetry of a cluster by having a smaller influx of Fz on the proximal side.

*γ* is the binding strength between pairs of proteins. Higher *γ* makes the stoichiometry between them stricter, lower *γ* makes them more uncorrelated. *λ* = 1 mimics wild-type whereas *λ* < 1 mimics the dsh[DIX] mutants. We assume that *λ* both affects the general influx of Dsh to the cluster and its binding with Fz.

*δ* represents the intracellular strength of mutual inhibition between Vang and Fz. Here, it scales linearly with the number of Fz or Vang monomers in the complex on the same cell side.

*ω* represents the preferred stoichiometry of Vang compared to Fmi, Fz, Dsh, and Pk. This parameter is conditioned on correct orientation of the cluster, meaning *ω* = 1 when the cluster lacks one of Fmi*_D_*, Fmi*_P_*, Fz*D*, Vang*_P_*, Dsh*_D_*, or Pk*_P_*, otherwise *ω* is larger than 1, thereby changing the preferred stoichiometry of Vang relative to the other proteins, e.g. by changing the conformation of the proteins and/or by phosphorylation of Vang. Effectively we are here suggesting that a currently unknown mechanism proofreads the completeness of the cluster before incorporation of additional Vang can occur.

In Fig. S10B, we analyze *N* = 50,000 clusters for *T* = 20,000. *α* = 0.968, *β* = 0.4, *γ* = 0.05, *δ* = 0.08, *ω* = 3, *λ* = 1 for wild type and *λ* = 0.2 for dsh[DIX]. This model captures the behavior observed in Figs. 1-6, and predicts that the amount of distal Vang anticorrelates with the amount of proximal Fmi and the amount of distal Fz. Conversely, in a *fz*^null^ background distal Vang correlates with proximal Fmi and distal Fz (Fig. S10C).

Nine cluster simulations are presented as movies illustrating examples of dynamic growth and decay trajectories taken by clusters (Movie S2).

### Supplementary figures

**Figure S1:**
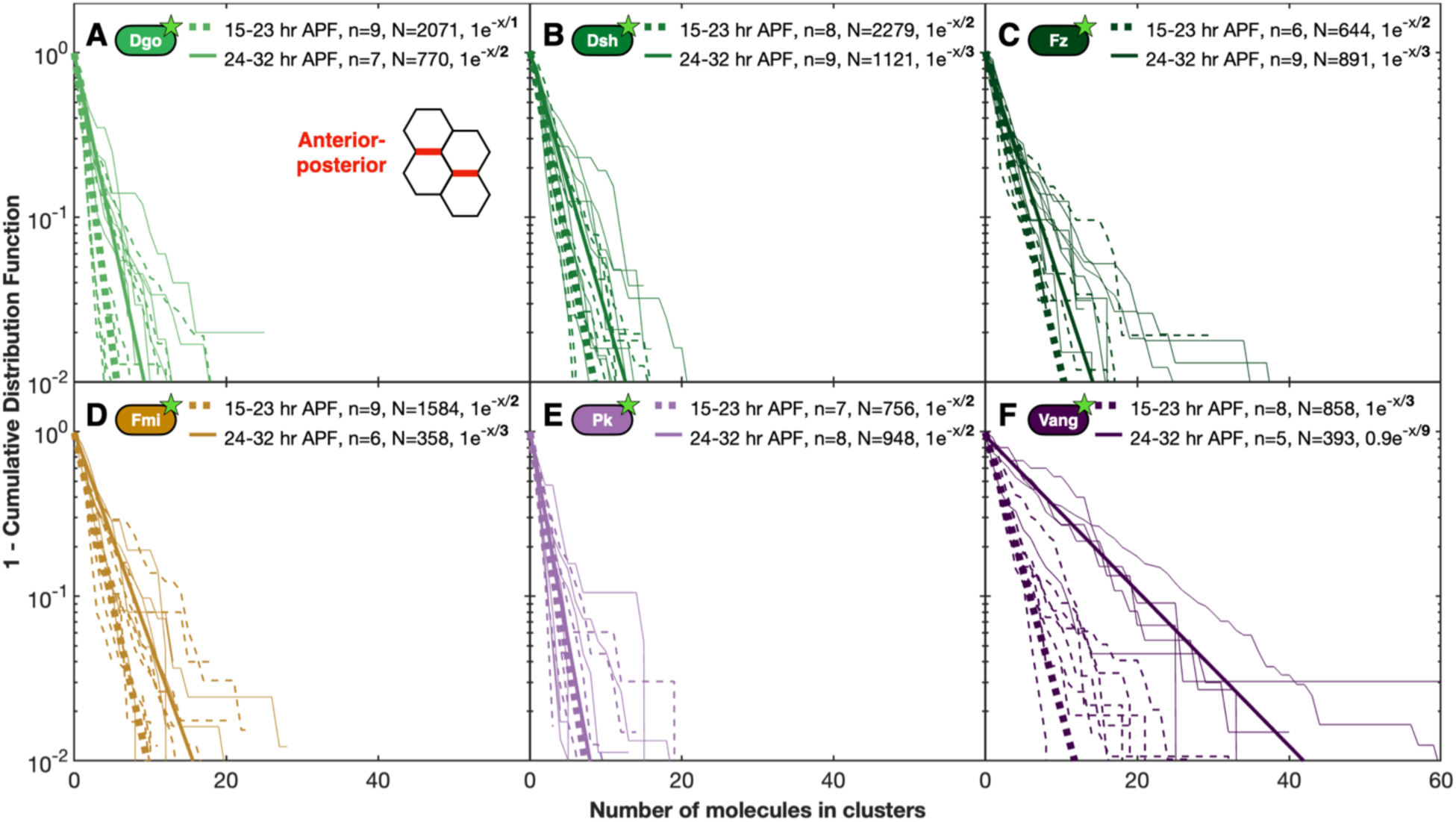
Cluster size distributions along anterior-posterior cell boundaries. **(A)** Dgo, **(B)** Dsh, **(C)** Fz, **(D)** Fmi, **(E)** Pk, and **(F)** Vang. Otherwise, similar to Figure 1 that shows the same along proximal-distal cell boundaries. When comparing to Figure 1, note, a difference in the scale on the *x*-axis.

**Figure S2:**
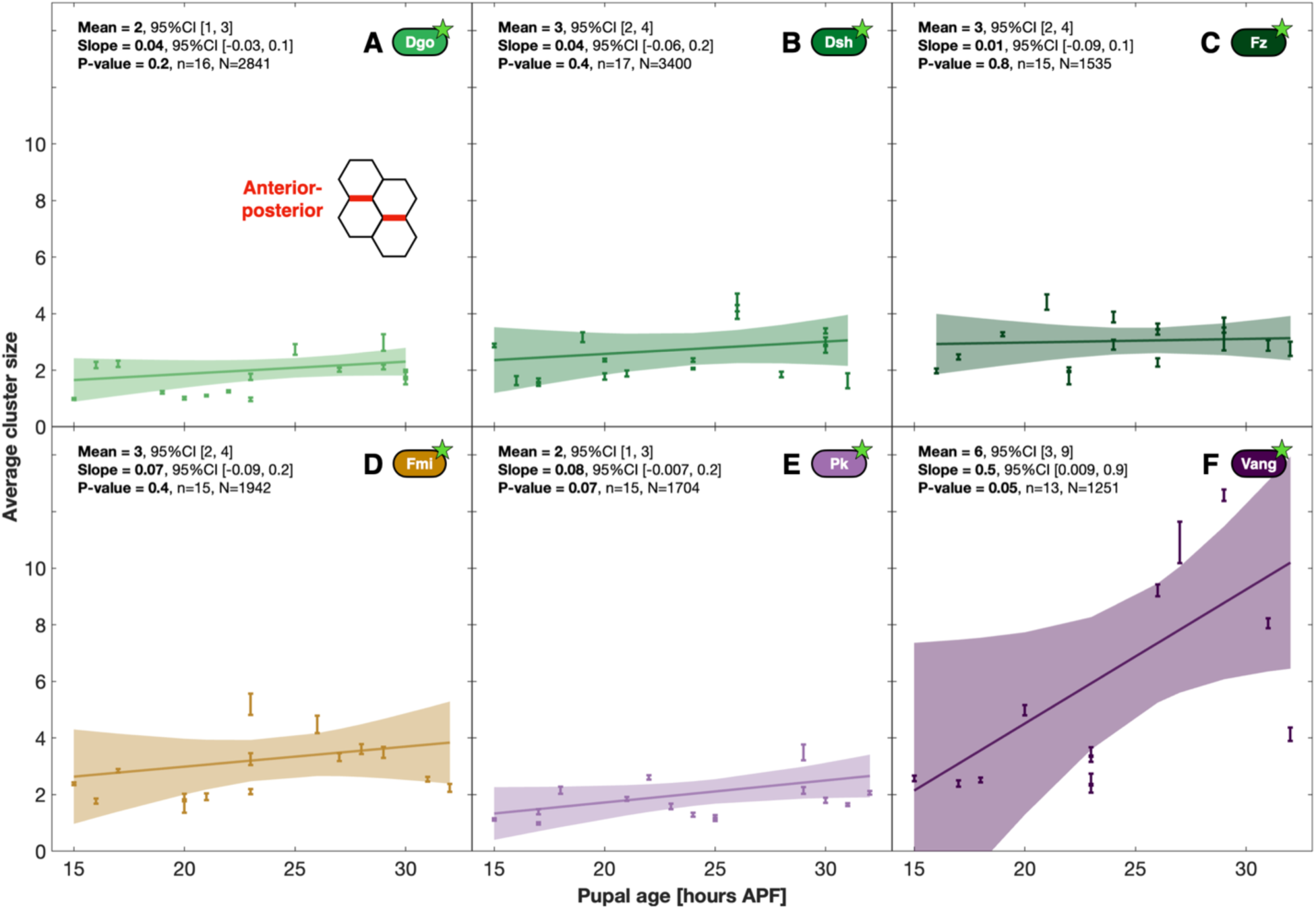
Planar cell polarity average cluster sizes along anterior-posterior (A-P) boundaries in uniformly labelled cells. **(A)** Diego (Dgo), **(B)** Dishevelled (Dsh), (**C)** Frizzled (Fz), **(D)** Flamingo (Fmi), **(E)** Prickle (Pk), and **(F)** Van Gogh (Vang). Dots, error bars, Mean, Slope, P-value, *n* and *N* as in Figure 2. Note, the small *y* scale relative to Figure 2. The underlying cluster size distributions for A-P boundaries are shown in Figure S1. We note that there exists no strict way of classifying clusters into boundaries.

**Figure S3:**
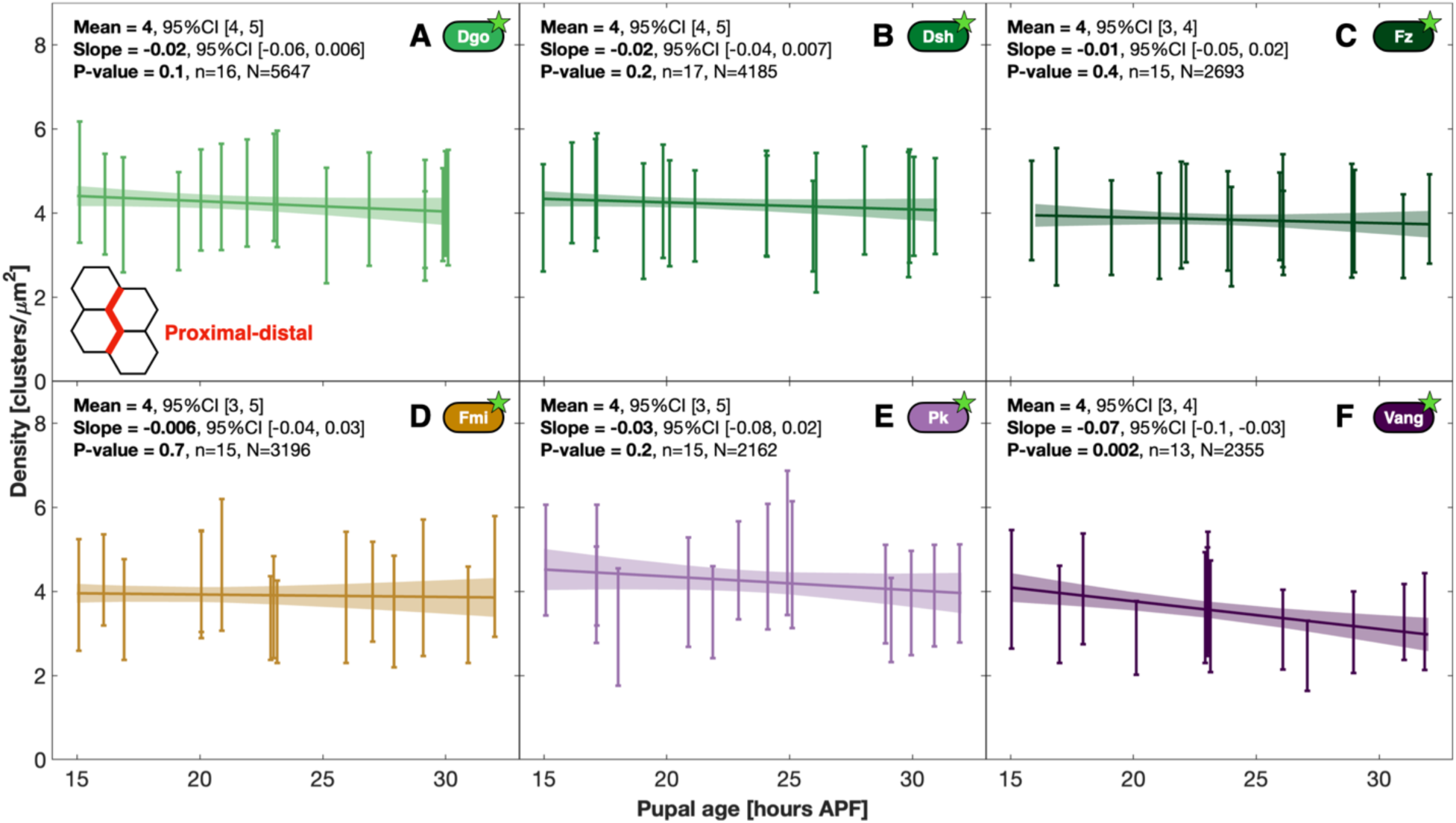
The number of clusters is roughly the same for all six core planar cell polarity (PCP) proteins and independent of the pupal age. For each identified cluster (see *Methods*), the number of clusters within a distance of √(1 μm^2^/π) is reported. Each error-bar represent one wing imaged and the standard deviation on the cluster density along proximal-distal boundaries in uniform expression of **(A)** Diego (Dgo), **(B)** Dishevelled (Dsh), **(C)** Frizzled (Fz), **(D)** Flamingo (Fmi), **(E)** Prickle (Pk), and **(F)** Van Gogh (Vang). A weighted linear regression model is shown through all samples for each condition. The mean denotes the predicted average density at 23.5h after puparium formation along with 95% confidence intervals (CI). The slope of the regression model is shown along with the P-value testing whether the density is depending on pupal age. *n* is the number independent samples imaged, and *N* is the total number of clusters.

**Figure S4:**
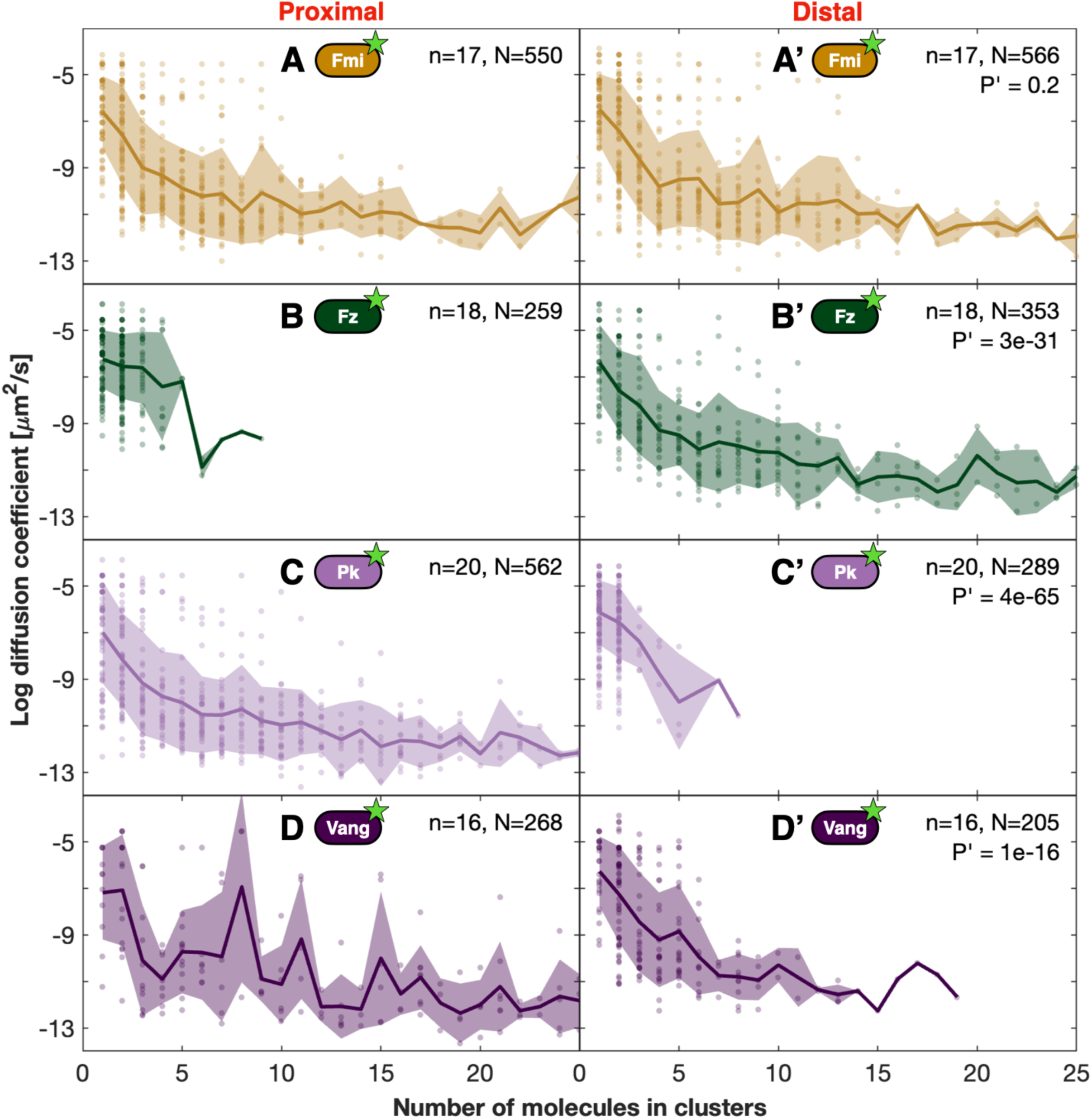
The diffusion coefficient, *D*, of subcomplexes as a function of cluster size in clone boundary experiments. **(A)** Proximal Flamingo (Fmi). **(A’)** Distal Fmi. **(B)** Proximal Frizzled (Fz). **(B’)** Distal Fz. **(C)** Proximal Prickle (Pk). **(C’)** Distal Pk. **(D)** Proximal Van Gogh (Vang). **(D’)** Distal Vang. The two-dimensional *D* is determined for each subcomplex identified by calculating *D = <r*^2^*>*/4*T,* where *<r*^2^*>* is the average squared displacement and *T* is the duration the subcomplex can be tracked. The P’ value tests whether *D* is different for identified proximal vs. distal clusters using a linear mixed-effects model test with model formula DiffCoef ∼ ClusterSize + P_or_D + ClusterSize * P_or_D. Each dot represents one (light colors) or more (darker colors) subcomplexes. The solid curve shows the mean *D f*or each cluster size and the standard deviation is shown in shaded colors. *n* denotes the number of wings imaged, and *<u>N</u>* the total number of subcomplexes tracked. Only subcomplexes with a track length of three or more frames are included.

**Figure S5:**
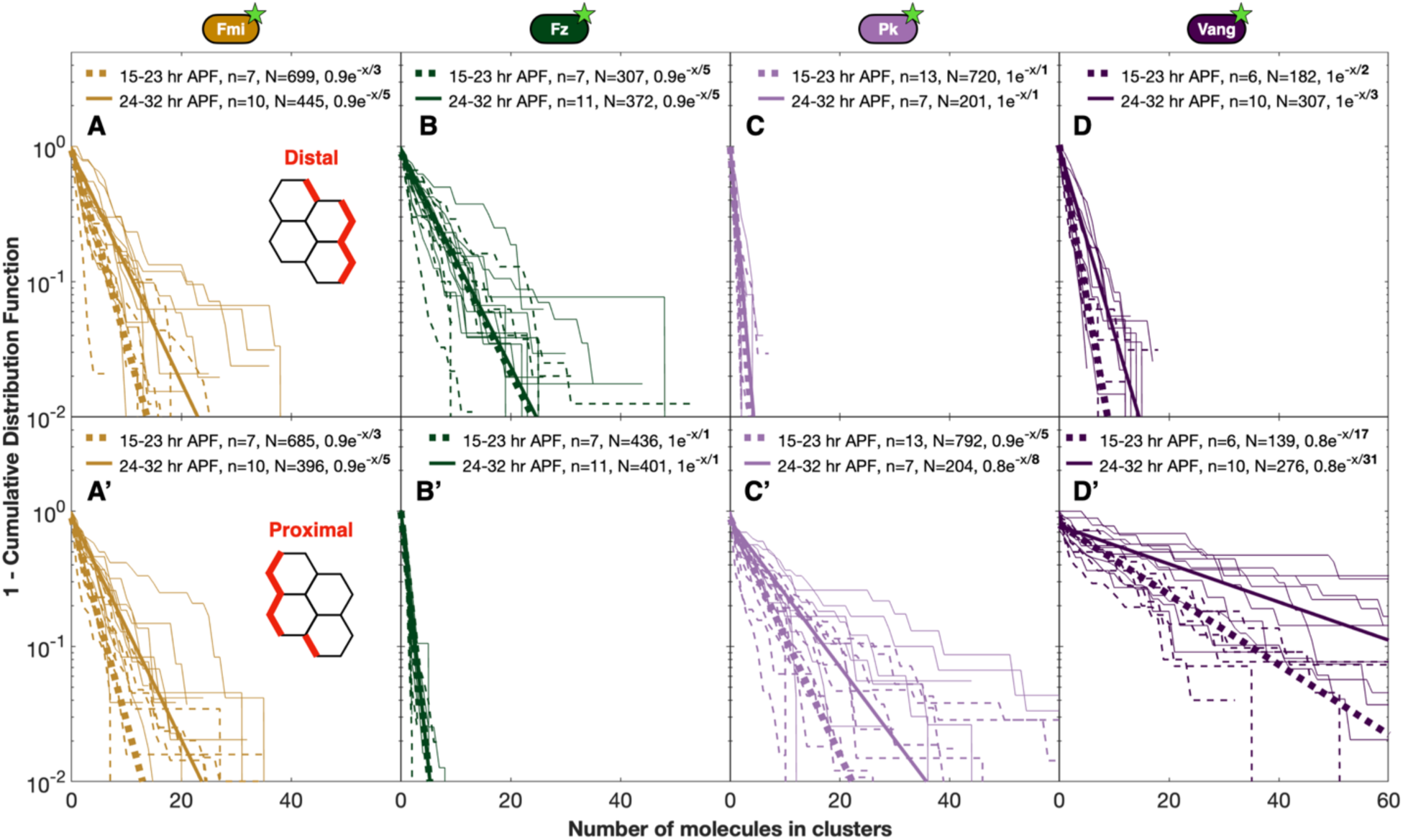
Cluster size distributions along proximal or distal cell boundaries. **(A)** Distal Fmi, **(B)** Distal Fz, **(C)** Distal Pk, and **(D)** Distal Vang. **(A’)** Proximal Fmi. **(B’)** Proximal Fz, **(C’)** Proximal Pk, and **(D’)** Proximal Vang. Data portrayed as in Figure 1. For age trends, see Figure 3. For anterior-posterior boundaries, see Figure S1.

**Figure S6:**
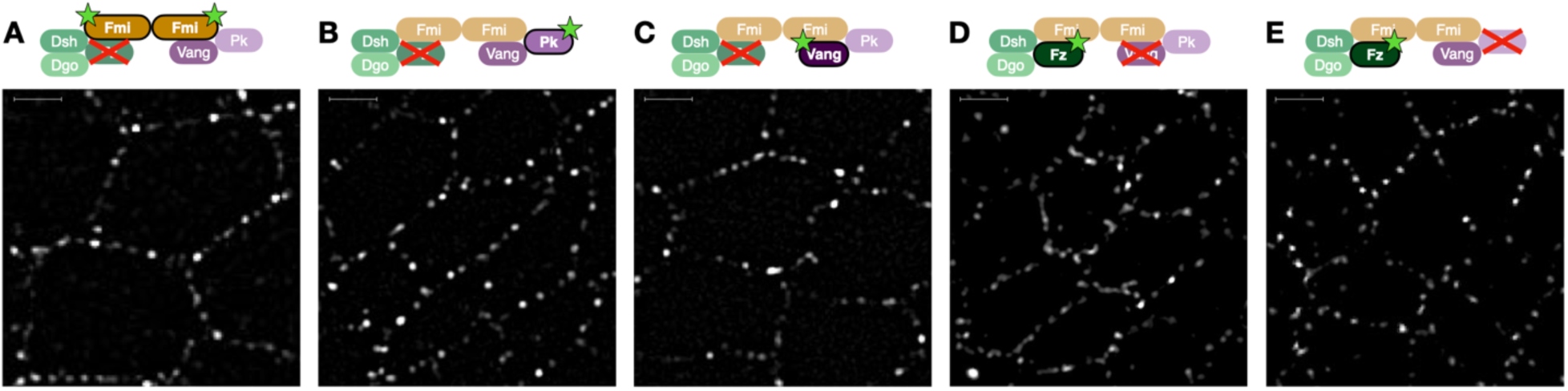
High-resolution sample images for loss-of-functions conditions in Figure 4. **(A)** Fmi::eGFP in *fz^R52^ / fz^R52^*, 17 hr APF, **(B)** Pk::eGFP in *fz^R52^ / fz^R52^*, 20 hr APF, **(C)** Vang::eGFP in *fz^R52^ / fz^R52^*, 32 hr APF, **(D)** Fz::eGFP in *vang^stbm6^ / vang^stbm6^*, 29 hr APF, **(E)** Fz::eGFP in *pk^pk-sple13^ / pk^pk-sple13^*, 18 hr APF, all acquired with TIRF Microscopy (see *Methods*). All scale bars are 2 μm. For quantification of cluster sizes along cell boundaries, see Figure 4.

**Figure S7.**
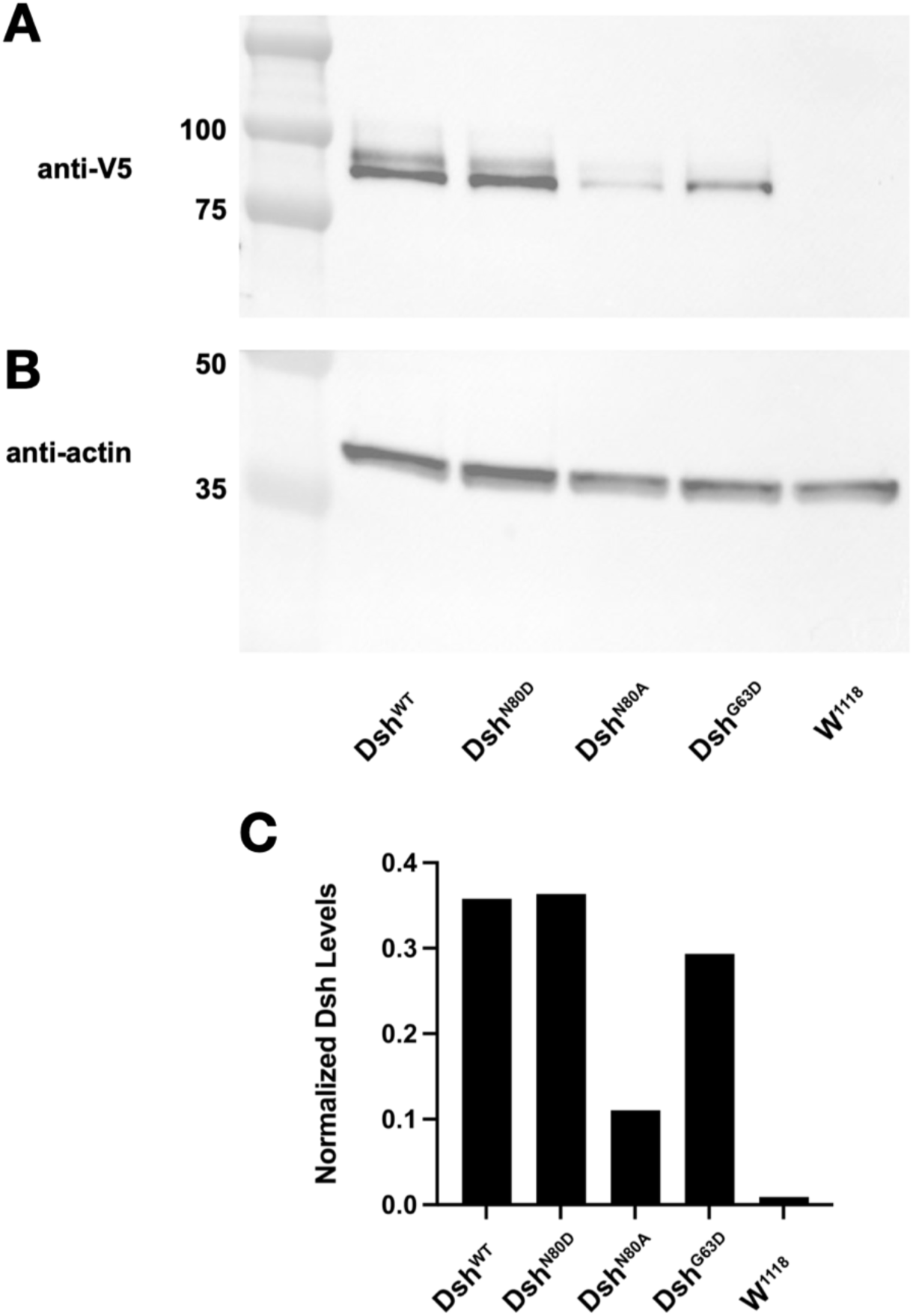
Dsh^DIX^ mutant protein levels by Western blot. Wing discs from female V5-tagged Dsh^WT^, Dsh^N80D^, Dsh^N80A^, Dsh^G63D^ and *w1118* larvae probed with **(A)** anti-V5 and **(B)** anti-actin. **(C)** Densitometric quantification of the Dsh protein levels normalized to actin level; values shown are the average of two independent experiments.

**Figure S8:**
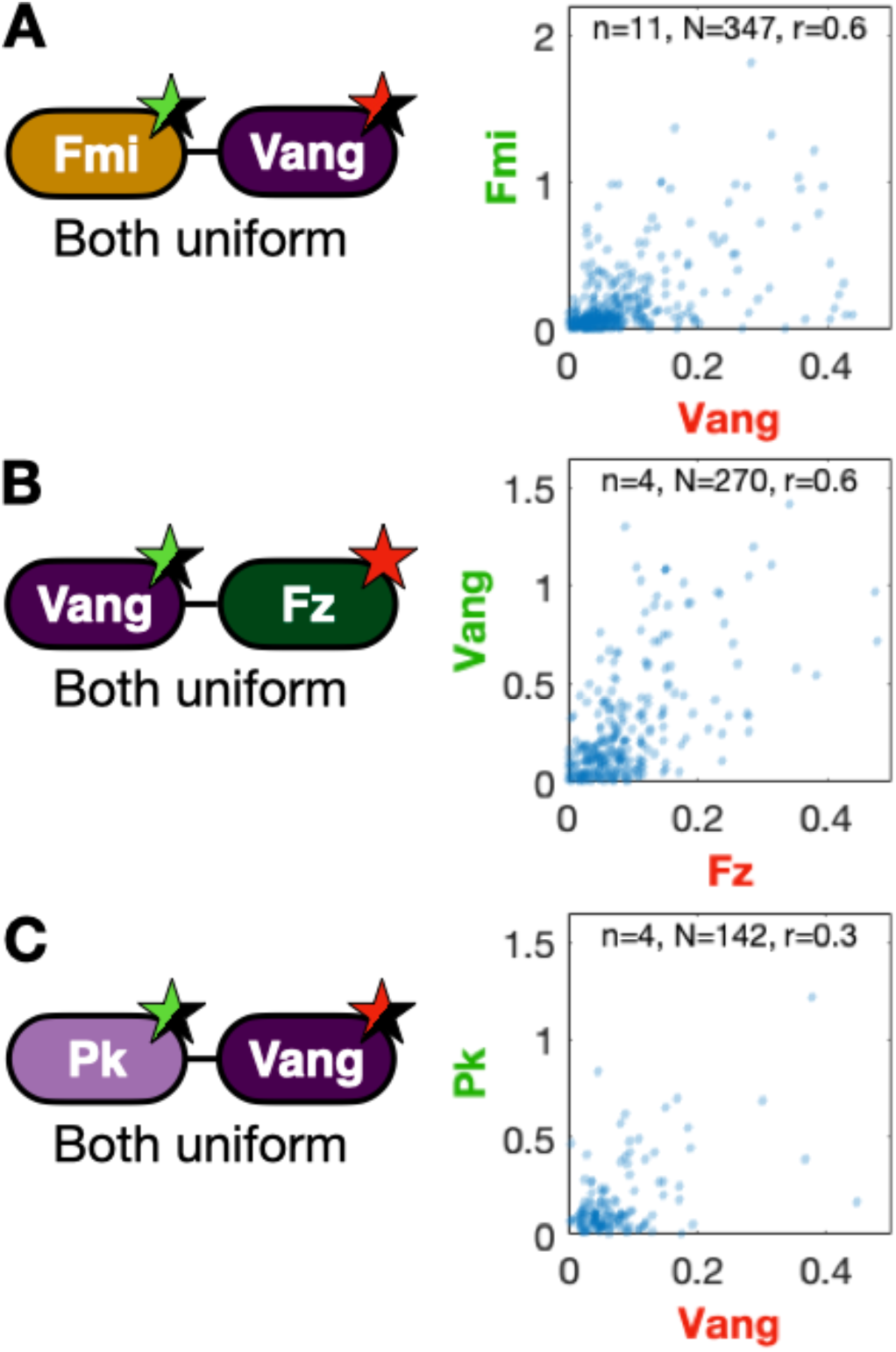
2-color correlations in uniform expression along anterior-posterior boundaries. **(A)** heterozygous Fmi::eGFP, heterozygous mScarlet::Vang, **(B)** heterozygous Vang::eGFP, homozygous Fz::mScarlet, and **(C)** heterozygous Pk::eGFP, heterozygous mScarlet::Vang. Otherwise, similar to Figure 6A-C which shows 2-color sample images and quantification along proximal-distal boundaries. Note, the smaller clusters along anterior-posterior boundaries vs. proximal-distal boundaries.

**Figure S9:**
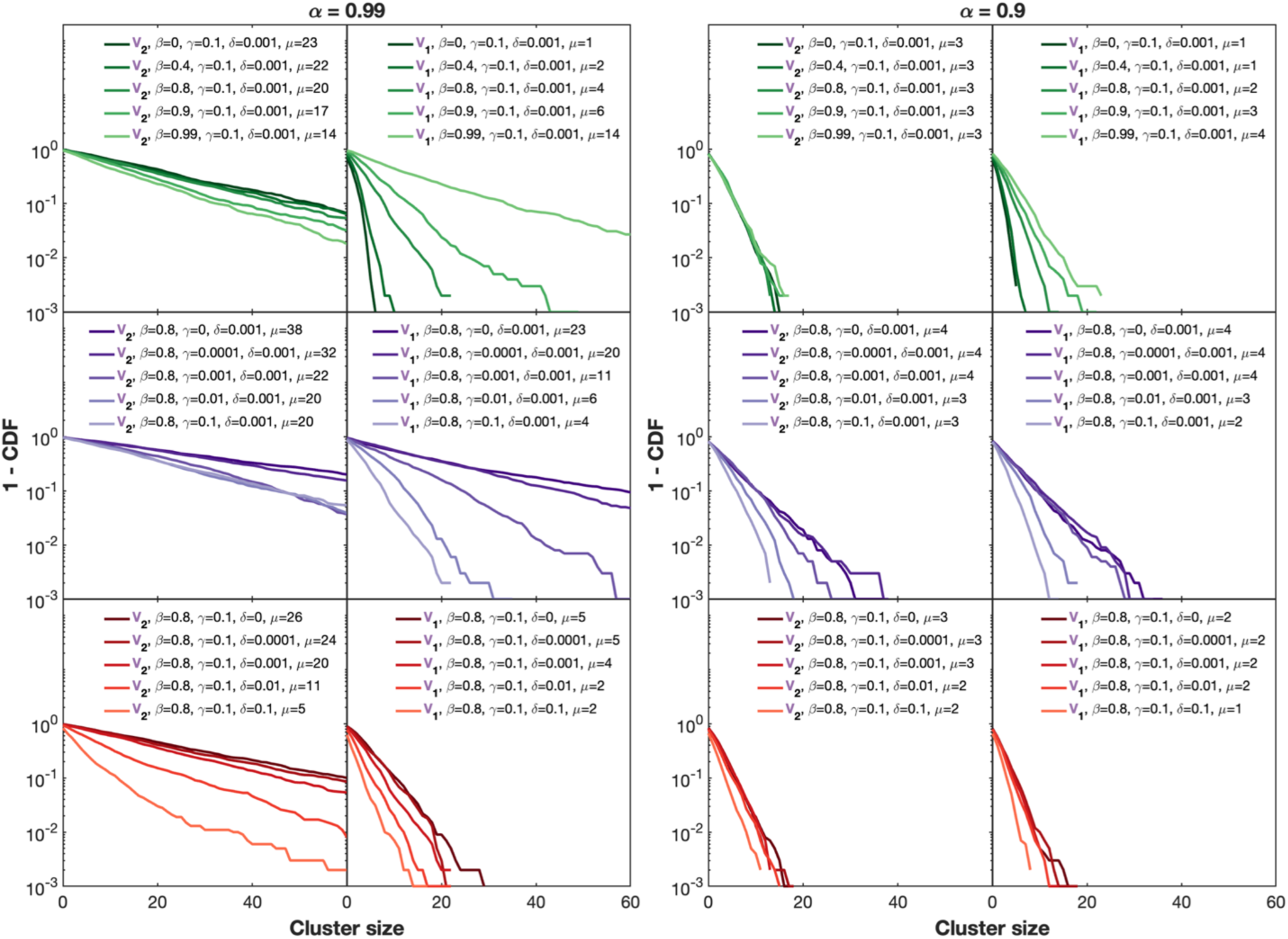
Simulated cluster size distributions across a set of parameters. Columns 1-2: For high *α* = 0.99, representing proximal-distal clusters at late pupal age. Columns 3-4: For slightly lower *α* = 0.9, representing anterior-posterior clusters or proximal-distal clusters at early pupal age. Row 1: Vary *β* for fixed *γ* and *δ*. Row 2: Vary gamma for fixed *β* and *δ*. Row 3: Vary *δ* for fixed beta and *γ*. Columns 1 and 3: Subcomplex *V* inside cell 2. Columns 2 and 4: Subcomplex *V* inside cell 1. *µ* denotes the average cluster size. Throughout, the number of simulated clusters *N* = 1000 are measured at the simulated time-point *T* = 10^6^ where the population of clusters has reached steady-state.

**Figure S10:**
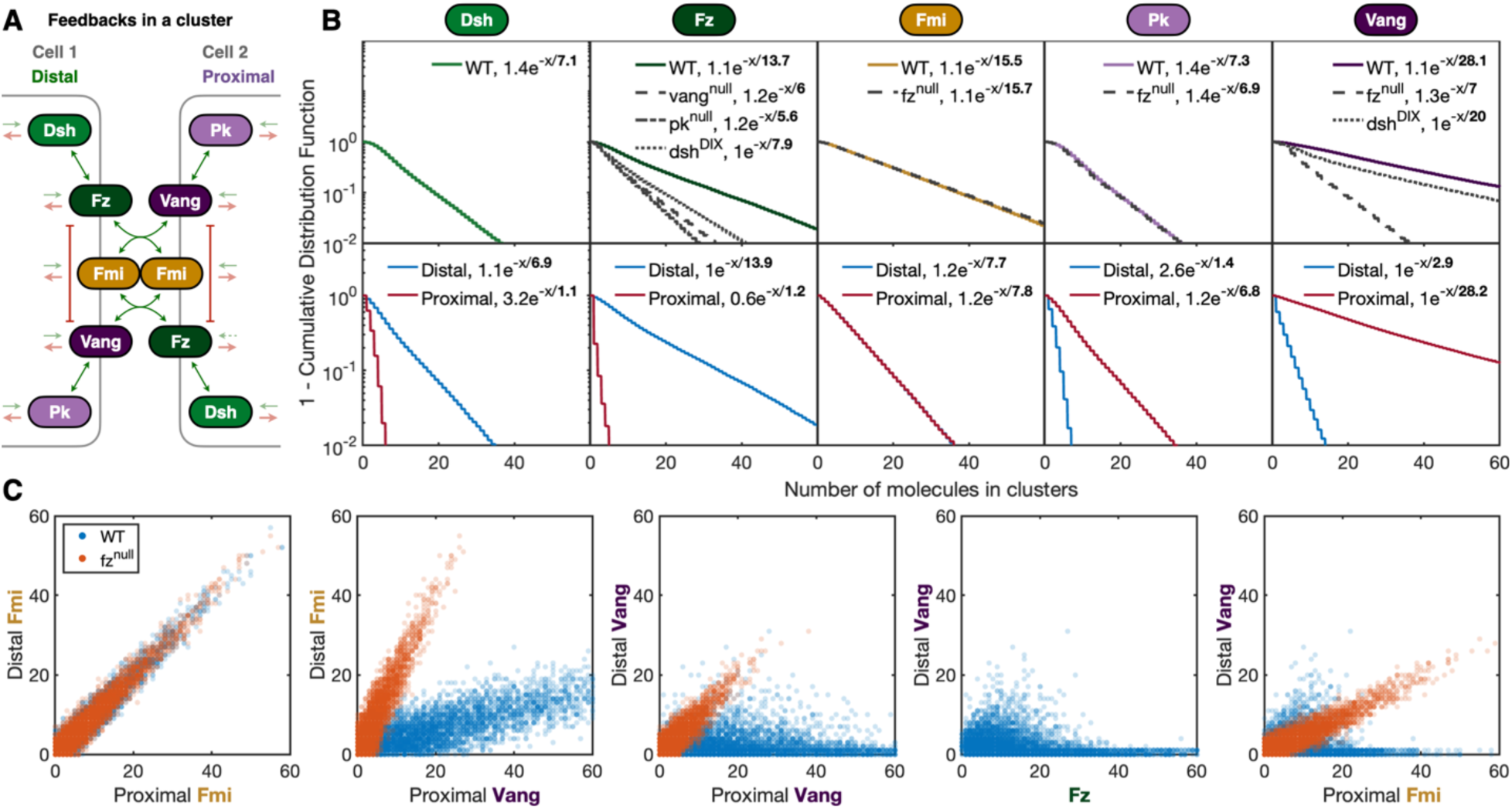
A five-component stochastic mathematical model of PCP cluster formation. **(A)** The model consists of Fmi, Fz, Vang, Dsh, and Pk. The proximal side is a mirror of the distal side, with the exception that the influx of Fz is less on the proximal side (dashed arrow). The Fz-Vang intercellular interaction is mediated through Fmi (see *Methods* and *Appendix* for details). **(B)** Cluster sizes in wild type (colored lines) summed across the boundary (top row, compare with Figures 1-2) and along proximal or distal boundaries (bottom row, compare with Figures 3 and S5) for *N =* 50,000 simulated clusters. Gray dashed lines indicate mutant simulations (compare with Figures 4-5). Note the *y*-axis is logarithmic, and the bold numbers indicate the mean number of monomers given by a single exponential fit. **(C)** Simulated two-color outcomes when looking at different pairs of proteins in wild type (blue) and pairs of proteins in a *fz^null^* background (red). Each dot represents one of the 50,000 simulated clusters.

**Figure S11:**
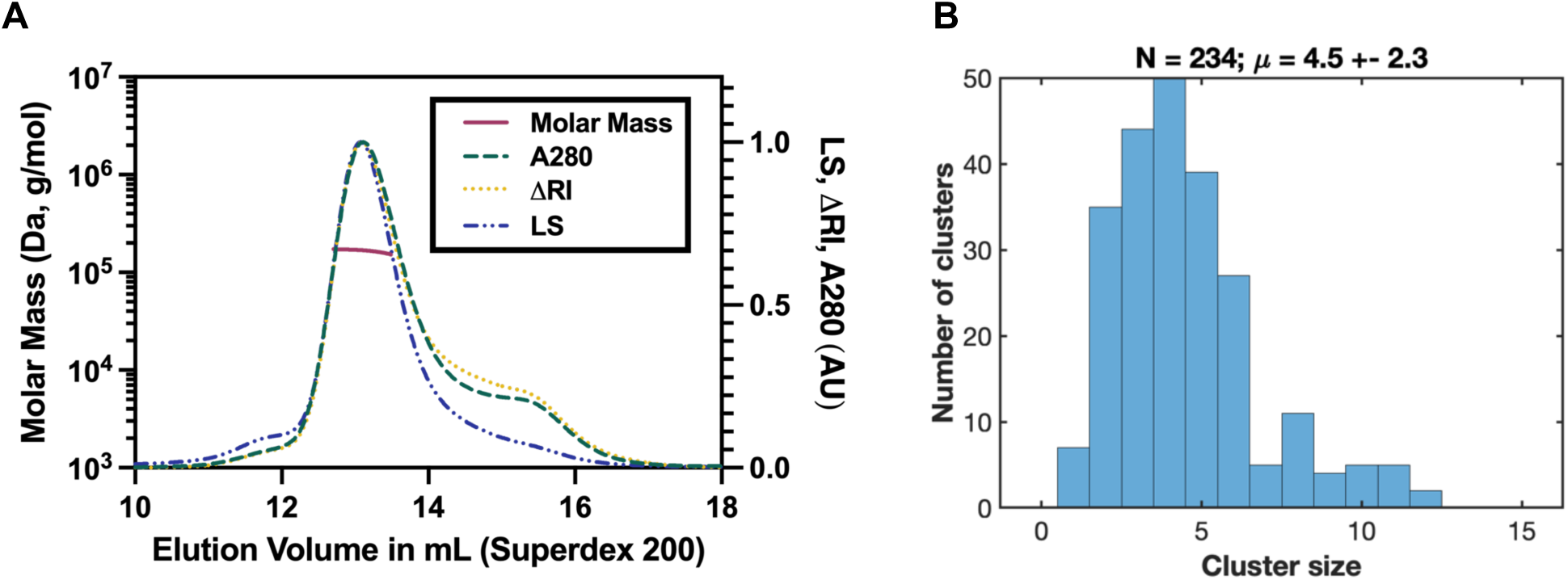
Validation of the counting method. **(A)** Molecular mass of Lyn-GFP-COMP pentamers was measured by Multi-Angle Light Scattering coupled to size-exclusion chromatography (SEC-MALS). Purified Lyn-GFP-COMP at 2mg/ml was run on a Superdex 200 10/300 column in line with multi-angle light scattering (MALS), absorption (A280) and refractive index (RI) detectors. The molar mass (left Y-axis) of Lyn-GFP-COMP was calculated over a range of elution volumes (X-axis), with a mean mass of 166.5kDa and median mass of 168.6kDa. Monomers of Lyn-GFP-COMP are predicted to be 34.2 kDa, so the measured masses are consistent with a pentamer with a predicted mass of 172.3 kDa. **(B)** Counting the purified pentameric GFP protein. Coverslips were washed with ethanol for 10 minutes, washed in water, flamed, and BSA added. A dilution of 10^-7^ of the SEC medium was used. The images were acquired on the same TIRF microscope and using the same settings as were used when imaging the wings, i.e., 50 ms exposure with 50% blue laser power. The bleaching traces were analyzed for all pixels using the same protocol as for the wings. The histogram includes data from three experiments. N is the total number of clusters identified, and μ is their average cluster size +/- their standard deviation.

**Figure S12:**
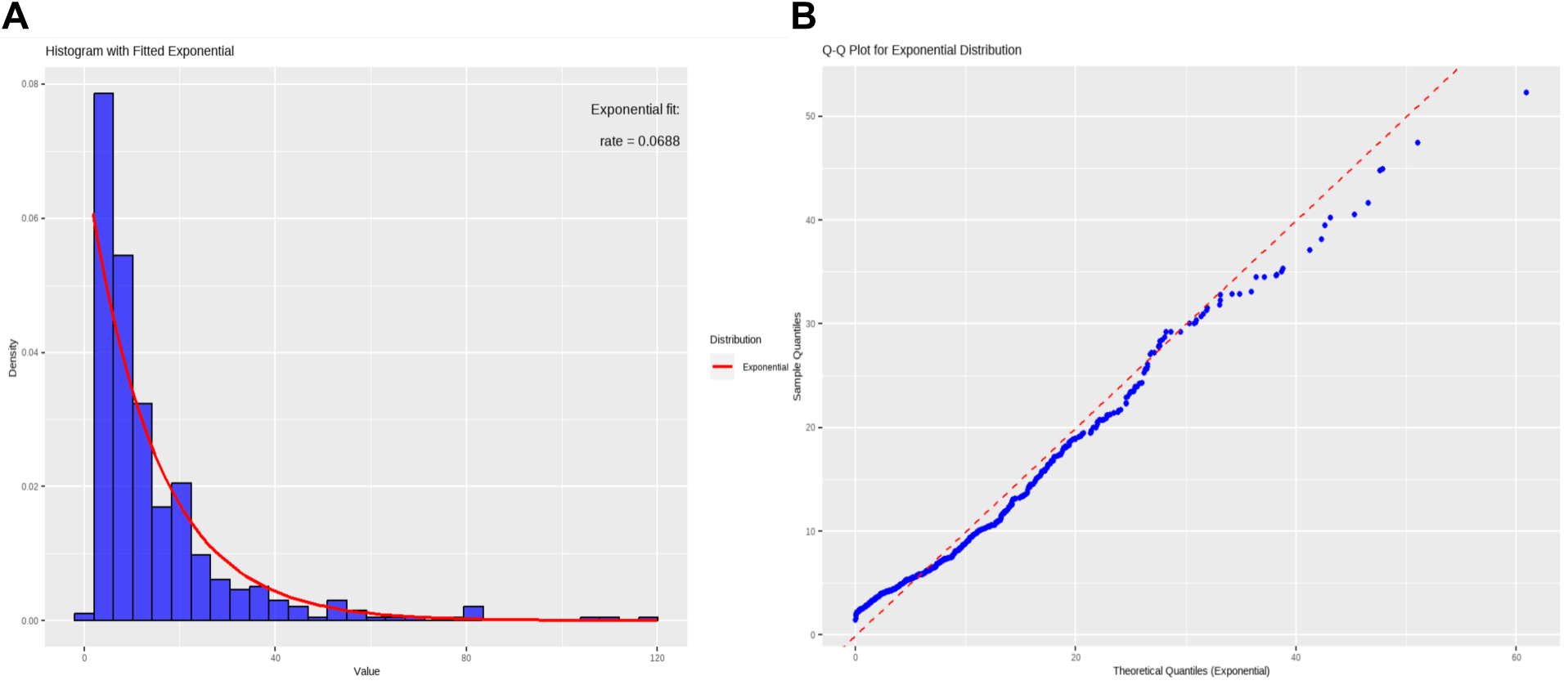
Visual tests of single exponential fits. **(A)** Histogram of Vang::eGFP cluster size in the age interval 15-23 hr APF shows that the data (blue bars) closely follows the exponential density plot (red curve; data from Figure 1). **(B)** Q-Q plot for Dgo::eGFP cluster size in the age interval 24-32 hr APF demonstrates that the data points align well with the reference diagonal line (shown in red; blue data points are from Figure 1).

**Movie S1:** 1.5 hour long time-lapse movie of a wing expressing Vang::eGFP starting at 17 hours after puparium formation. 10 frames shown per second with each frame corresponding to 10 seconds. Acquired on a Total Internal Reflection Fluorescence microscope (see *Methods*) with the blue laser repeatedly set at 1% laser power for an exposure of 50 ms followed by turning off the laser for 9950 ms. Distal is to the right. Note the roughly stable position of clusters and a few cell divisions. It is likely that very small clusters are not detected in this movie due to the low laser power.

**Movie S2:** Mathematical model simulation of planar cell polarity (PCP) cluster assembly. Nine panels are shown illustrating the evolution of nine independent clusters. Each colored thin horizontal bar represents one monomer of a protein. The number of monomers of each protein can be read on the *y*-axis. Inside each panel, the first three columns represent cell 1 and the last three columns represent the neighboring cell 2. Positive monomers (assembling upwards) represent monomers on the ‘correct’ side and negative monomers (assembling downwards) represent monomers on the ‘incorrect’ side. The first column shows proximal Dsh (light green), and distal Pk (light purple). The second column shows proximal Fz (dark green), and distal Vang (dark purple). The third column shows proximal Fmi (orange). The fourth column shows distal Fmi (orange). The fifth column shows proximal Vang (dark purple) and distal Fz (dark green). Finally, the sixth column shows proximal Pk (light purple) and distal Dsh (light green). Notice how small clusters can be oriented incorrectly whereas large clusters will be oriented correctly and that the degree of asymmetry increases with cluster size. At any given time point, this simulation will give rise to single exponential cluster size distributions for each of the core proteins. For details see ‘The more complete math model’ in the Appendix.

